# Balance Between EGF/STAT1 and IFNγ/NRF2 Signaling Controls ECM1 Expression and Determines Hepatic Homeostasis versus Chronic Liver Disease

**DOI:** 10.1101/2024.02.17.580798

**Authors:** Yujia Li, Chenjun Huang, Weiguo Fan, Seddik Hammad, Lea Berger, Cyrill Géraud, Ye Yao, Frederik Link, Chenhao Tong, Zeribe C. Nwosu, Pia Erdösi, Weronika Piorońska, Kerry Gould, Christoph Meyer, Rilu Feng, Hui Liu, Chen Shao, Bing Sun, Huiguo Ding, Roman Liebe, Matthias P. A. Ebert, Honglei Weng, Chunfang Gao, Peter ten Dijke, Steven Dooley, Sai Wang

## Abstract

In healthy livers, extracellular matrix protein 1 (ECM1) is essential for liver homeostasis by keeping latent transforming growth factor-β (LTGF-β) quiescent. Upon hepatocyte damage, ECM1 is downregulated, facilitating LTGF-β activation and fibrogenesis. However, little is known about how hepatic ECM1 is regulated. Here we found in healthy hepatocytes, EGF/EGFR signaling sustains ECM1 expression through phosphorylating STAT1 at S727, enhancing its binding to the *ECM1* promoter and boosting gene transcription. During liver inflammation, accumulating IFNγ disrupts this process by downregulating EGFR and inhibiting EGF/EGFR/STAT1-mediated *ECM1* promoter binding. Mechanistically, IFNγ-induced STAT1 phosphorylation at Y701 impairs the binding of p-STAT1 S727 to the ECM1 promoter. Additionally, IFNγ induces NRF2 nuclear translocation, which repressively binds to the *ECM1* promoter, further reducing its expression. These findings were confirmed in several chronic liver disease (CLD) mouse models. Moreover, AAV8-ECM1 significantly attenuates liver fibrosis and injuries in Western diet (WD)-fed mice. Notably, in patients with CLD, ECM1 levels align with EGFR expression, while NRF2 and LTGF-β activation show a negative correlation with both.

## Introduction

Extracellular matrix protein 1 (ECM1) is a secreted 85-kDa glycoprotein (Mathieu *et al*, 1994) primarily found in the epidermis and dermis (Smits *et al*, 2000), where it functions to maintain the integrity and homeostasis of the skin (Sercu *et al*, 2009). ECM1 is also involved in the development of cancer and its levels are elevated in most malignant epithelial and metastatic tumours, including breast cancer, colon cancer, ovarian cancer, and melanoma (Sercu *et al*, 2008; Wang *et al*, 2003). In contrast, in our previous studies, we identified ECM1 as gatekeeper of liver homeostasis by interacting with and inhibiting αv integrin-, thrombospondin 1 (TSP-1)-, ADAMTS protease 1 (ADAMTS1)-, and matrix metalloproteinase (MMP)-mediated latent TGF-β (LTGF-β) activation (Fan *et al*, 2019; Li *et al*, 2022; Link *et al*, 2024). Mice lacking *Ecm1* gene display severe hepatic fibrosis and die between ages of 8 to 12 weeks (Fan *et al*., 2019). ECM1 is produced mainly by hepatocytes and quiescent hepatic stellate cells (HSCs), deposited in the extracellular matrix for its function (Fan *et al*., 2019; Filliol *et al*, 2022). Importantly, ECM1 expression is significantly downregulated following hepatocyte injury and HSC activation (Fan *et al*., 2019). However, since hepatocytes are the most vulnerable cells in the liver (Gribben *et al*, 2024), the reduction of ECM1 mediated by damaged hepatocytes, especially at the early stages of chronic liver disease (CLD) prior to HSC activation, is considered a key factor in initiating further liver injuries (Fan *et al*., 2019).

Several tested mouse models of liver disease e.g., carbon tetrachloride (CCl_4_)-, metabolic dysfunction-associated steatohepatitis (MASH)-, and bile duct ligation (BDL)-associated liver injury consistently display Ecm1 downregulation (Fan *et al*., 2019), suggesting a common initiating mechanism of liver fibrogenesis. A progressive loss of ECM1 with disease severity is present in patients with liver fibrosis/cirrhosis (Fan *et al*., 2019), and this also correlates with poorer prognosis in hepatocellular carcinoma (HCC) patients (Bai *et al*, 2020; Gao *et al*, 2014). It is therefore of great interest to investigate ECM1 expression regulation in the liver context. Our research focused on the regulation of ECM1 expression in hepatocytes, as they are one of the main producers and the downregulation of ECM1 by injured hepatocytes in the early stages of CLD is a crucial factor in driving disease progression.

## Results

### EGF promotes ECM1 expression in hepatocytes

As reported previously, hepatic ECM1 expression undergoes a significant downregulation in response to liver injury (Fan *et al*., 2019). To gain a comprehensive understanding of its modulation in liver pathology, we analyzed the *ECM1* mRNA expression in liver tissue of CLD patients sourced from the Gene Expression Omnibus (GEO) online database. It revealed a consistent reduction in *ECM1* expression across various cohorts of CLD patients, demonstrating a correlation with disease progression **(Figure 1A)**. Specifically, this decline was evident in patients with metabolic dysfunction-associated steatotic liver disease (MASLD, n=15) and metabolic dysfunction-associated steatohepatitis (MASH, n=16), in comparison to healthy individuals with normal weight (n=14) and obesity (n=12) (Suppli *et al*, 2019) **(Figure 1A, left)**. Furthermore, *ECM1* expression was notably diminished in alcoholic cirrhosis patients (n=67) relative to those with non-severe alcoholic hepatitis (n=13) and alcoholic steatosis (n=6) (Trepo *et al*, 2018) **(Figure 1A, middle)**. In addition, the observed significant decrease in *ECM1* mRNA expression extended to tumor tissue from HBV-associated HCC patients (n=21) compared to non-neoplastic liver tissues (Yoo *et al*, 2017) **(Figure 1A, right)**. This consistent pattern underscores the potential role of ECM1 in the pathogenesis and progression of various liver diseases.

**Figure 1.**
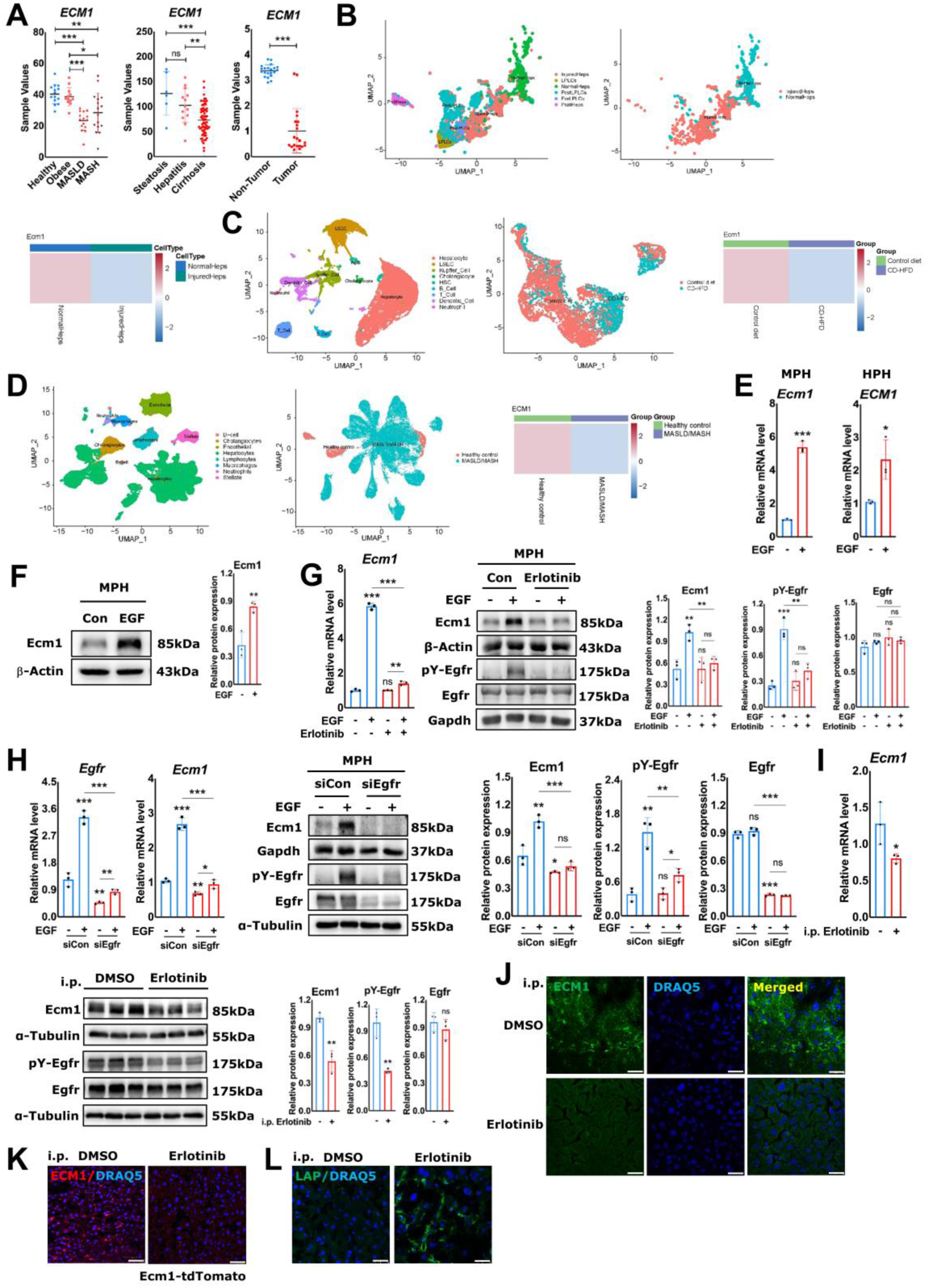
EGF-EGFR signaling maintains Ecm1 expression in hepatocytes of quiescent liver. **(A)** mRNA expression of hepatic *ECM1* in CLD patients, extracted from the GEO Dataset GSE126848, GSE103580 and GSE94660, respectively. UMAP visualization of cell populations and hepatocytes, and heatmap of *ECM1* mRNA expression in **(B)** DDC mouse model (GSE193850), **(C)** CD-HFD fed mice (GSE232182), and **(D)** patients with CLD (GSE202379). **(E)** qRT-PCR for *ECM1* mRNA expression in MPHs and HPHs with or without EGF treatment. **(F)** Immunoblotting of Ecm1 in MPHs treated with EGF. **(G)** qRT-PCR and immunoblotting showing the effect of erlotinib on Ecm1 expression and Egfr Y1068 phosphorylation in MPHs with or without EGF treatment. **(H)** Effects of *Egfr* knockdown on mRNA and protein expression of Ecm1 and Egfr in EGF-treated MPHs by qRT-PCR and immunoblotting. **(I)** qRT-PCR and immunoblotting for Ecm1 in liver tissues from DMSO or erlotinib-treated mice. DMSO was administered as a placebo. **(J)** Representative IF staining for Ecm1 expression in liver tissues from DMSO or erlotinib-treated mice. DRAQ5 fluorescent probe stains DNA. **(K)** Representative tdTomato fluorescence of liver tissues from Ecm1-tdTomato mice treated with DMSO or erlotinib. **(L)** Representative IF staining for LAP-D R58 in liver tissues from DMSO or erlotinib-treated mice. Scale bar, 25µm. The results of qRT-PCR were normalized to *PPIA*. In immunoblotting, β-Actin, Gapdh and α-Tubulin are loading controls. Quantification of protein expression was performed by ImageJ (National Institutes of Health, Bethesda, Maryland, USA). *P*-values were calculated by unpaired Student’s t test. Bars represent the mean ± SD. *, *P*<0.05; **, *P*<0.01; ***, *P*<0.001.

Our previous study demonstrated that healthy hepatocytes are one of the primary producers of ECM1 in the liver, as identified through qRT-PCR and immunoblotting (Fan *et al*., 2019). To further confirm the specific role of hepatocytes in maintaining ECM1 expression, we analyzed several scRNA-seq datasets from various mouse models of liver injury and patients with CLD. In 3,5-diethoxycarbonyl-1,4-dihydrocollidine (DDC) diet-injured mouse model (GSE193850) (Li *et al*, 2023), we classified the cell clusters into diverse categories **(Figure 1B, first panel)** based on the marker genes **(Suppl. Fig. 1A)**, among others, InjuredHeps and NormalHeps were selected for further analysis **(Figure 1B, second panel)**. Heatmap showed that in the injured hepatocytes, the expression of *Ecm1* was significantly deseased as compared to normal hepatocytes **(Figure 1B, third panel)**. Similarly, *Ecm1* expression in hepatocytes was also reduced in choline-deficient high-fat diet (CD-HFD)-fed mice (GSE232182) compared to the control diet **(Suppl. Fig. 1B; Figure 1C)**. Furthermore, in CLD patients (GSE202379) (Gribben *et al*., 2024), as the MASLD/MASH developed, the expression of *ECM1* in hepatocytes likewise progressively declined relative to healthy donors **(Suppl. Fig. 1C; Figure 1D)**. Since hepatocytes are the cell type most affected by the disease and exhibit a significant downregulation of ECM1 expression before HSC activation upon liver injury (Fan *et al*., 2019; Gribben *et al*., 2024), this suggests that the reduction in ECM1 by injured hepatocytes may be a key factor contributing to CLD progression. Therefore, our study focuses on investigating the mechanisms underlying ECM1 regulation in hepatocytes.

To get a broader understanding of signaling pathways and transcription factors potentially implicated in the regulation of *Ecm1* transcription, we performed an *in silico* analysis of the *Ecm1* gene (NCBI Gene ID: 13601) promoter (−2000bp ∼ +200bp relative to transcription start site (TSS)) using PROMO (https://alggen.lsi.upc.es/cgi-bin/promo_v3/promo/promoinit.cgi?dirDB=TF_8.3), GeneCards (ECM1 Gene - GeneCards) and JASPAR (https://jaspar.genereg.net/), and identified multiple binding sites for candidate transcription factors, including, among others, Fos, Jun, cMyc, NF-κB, and Stat1 **(Suppl. Fig. 2)**. NF-κB is mainly regulated by inflammatory cell signaling induced by TNF-α and LPS. Fos, Jun, cMyc, and Stat1 are downstream transcription factors of growth factors, especially EGF and HGF. Additionally, Stat1 is a major downstream mediator of the interferon family. As these signaling molecules all are having prominent roles in liver physiology and pathophysiology, they are promising candidates as crucial regulators of hepatic Ecm1 expression.

Next, we tested previously described pathways for their impact on *Ecm1* mRNA expression levels. 4ng/ml TNF-α or 5µg/ml LPS treatment did not impact on *Ecm1* mRNA expression in mouse primary hepatocytes (MPHs) **(Suppl. Fig. 3)**; EGF and HGF treatment consistently induced *ECM1* expression on mRNA levels in MPHs, human primary hepatocytes (HPHs) and AML12 cells **(Figure 1E**, **Suppl. Fig. 4A)**, as well as promoted Ecm1 protein expression in MPHs **(Figure 1F)** and AML12 cells **(Suppl. Fig. 4B)**.

### EGF/EGFR signaling contributes to physiological Ecm1 expression in liver homeostasis

Given the crucial role of the EGFR in the EGF signaling pathway (Zeineldin & Hudson, 2006), we applied erlotinib, a selective EGFR inhibitor in EGF-treated AML12 cells and MPHs. qRT-PCR and Western blotting analysis confirmed that erlotinib substantially inhibited EGF-induced Egfr phosphorylation and Ecm1 expression in both cell systems **(Figure 1G; Suppl. Fig. 4C)**. In line, siRNA mediated *Egfr* knockdown as well suppressed Egfr expression/phosphorylation levels, and Ecm1 expression in MPHs upon treatment with EGF **(Figure 1H)**. To confirm our findings *in vivo*, EGF was injected to WT mice through tail vein (200μg/kg body weight (BW)). Results showed that EGF treatment did not increase Ecm1 expression *in vivo*, which might be due to the physiological level of EGF in the healthy liver **(Suppl. Fig. 5)**. Alternatively, we applied erlotinib (40mg/kg BW, for 2 days) to WT mice intraperitoneally (i.p.) and analysed the liver tissue 48hrs later. qRT-PCR, Western blotting, and immunofluorescence (IF) analysis showed a significant decrease in Ecm1 expression in the erlotinib-treated group compared to the control group **(Figure 1I, J)**. Further evidence of this downregulation was demonstrated in Ecm1-tdTomato transgenic mice. In the liver tissue of these mice, Ecm1 was visualized by tdTomato fluorescence, which showed a reduction in red florescence signal under the same erlotinib treatment conditions **(Figure 1K)**. Consistent with our previous finding that ECM1 loss leads to the activation of latent TGF-β1 (LTGF-β1), erlotinib injection significantly activated LTGF-β1, as indicated by the elevated expression of LAP-D R58, a breakdown product of the TGF-β1 latency-associated peptide (LAP) that is detectable in the ECM upon LTGF-β1 activation (**Figure 1L**).

These results suggest that constant/basal EGF-EGFR signaling maintains Ecm1 expression in healthy livers.

### EGF induces Ecm1 expression through Stat1

To elucidate potential downstream components of the EGF-EGFR signaling pathway that regulate Ecm1 expression, we tested several transcription factors as predicted from the *in silico* promoter study, such as Fos, Jun, cMyc, and Stat1, together with its major downstream Erk signaling, by depleting their availability with siRNA interference experiments. Knockdown of Fos, Jun, or cMyc, Erk1 and Erk2 did not interfere with EGF-induced Ecm1 expression **(Suppl. Fig. 6A-E),** whereas qRT-PCR and immunoblot data revealed that *Stat1* silencing by siRNA markedly suppressed EGF-induced Ecm1 expression in MPHs **(Figure 2A)** and AML12 cells **(Suppl. Fig. 7A)**. This suggested Stat1 as a promising candidate. In line, 1hr EGF treatment of MPHs induced phosphorylation of Stat1 at S727 phosphorylation site, which remained stable for at least 24hrs **(Figure 2B)**. As Stat1 is also an EGF-EGFR downstream target (Wang, 2017), its total expression and phosphorylation at S727 were measured in EGF-treated MPHs with or without siRNA targeting *Egfr*. Western blotting showed EGF-induced total expression and phosphorylation of Stat1 to be dependent on Egfr **(Figure 2C)**. Notably, Egfr inactivation also inhibited the phosphorylation of Stat1 at position S727 in the mice treated with erlotinib **(Figure 2D)**. Preliminary data on HGF signaling, showing that depleting Stat1 did not abrogate Ecm1 expression induction in MPHs, suggest involvement of alternate pathways, whose detailed delineation is subject matter of an ongoing investigation **(Suppl. Fig. 8)**.

**Figure 2.**
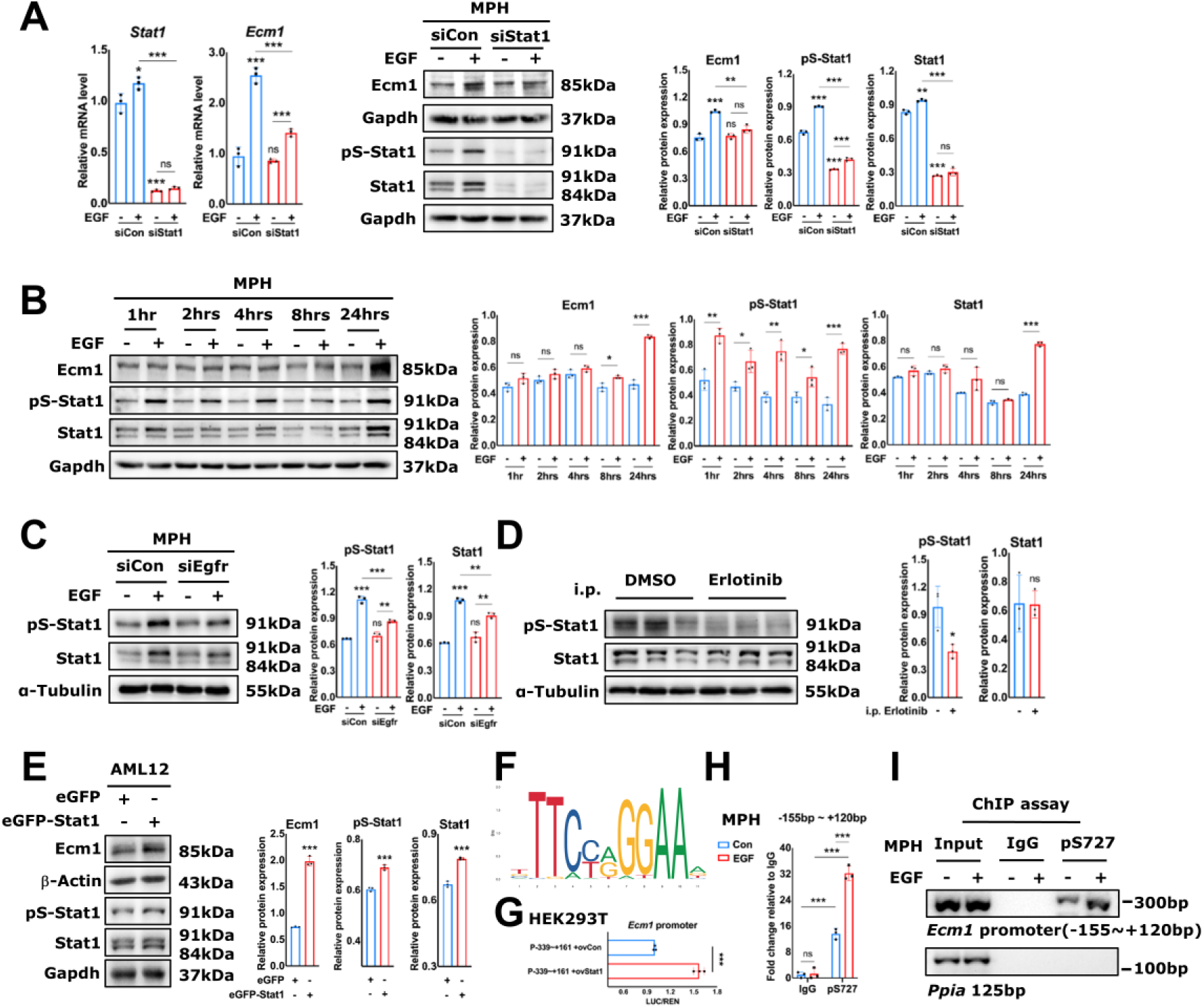
EGF regulates homeostatic Ecm1 expression in hepatocytes through Stat1. **(A)** qRT-PCR and immunoblotting of Ecm1 and Stat1 in EGF-treated MPHs with or without *Stat1* knockdown. **(B)** Immunoblotting showing the effect of EGF on target proteins in MPHs at different time points. **(C)** Immunoblotting for p-Stat1 S727 and Stat1 in EGF-treated MPHs with or without knockdown *Egfr*. **(D)** Immunoblotting for p-Stat1 S727 and Stat1 in liver tissue from DMSO or erlotinib-treated mice. DMSO was administered as a placebo. **(E)** Immunoblotting for Ecm1, p-Stat1 S727 and total Stat1 in AML12 cells transfected with expression vectors. **(F)** Predicted binding motif of Stat1 to *Ecm1* promoter by Jaspar. **(G)** Luciferase reporter assay analyzing the activity of *Ecm1* promoter (−339 ∼ +161bp) in HEK293T cells transfected with or without Stat1 plasmid. **(H)** ChIP qRT-PCR showing the binding of S727 phosphorylated Stat1 to the *Ecm1* gene promoter in MPHs, with or without EGF treatment. Fragment “-155bp ∼ +120bp’’ is calculated relative to the *Ecm1* gene TSS. Rabbit IgG-bound chromatin served as negative control. **(I)** The amplified products of ChIP PCR are shown as bands on 2% agarose gel. *Ppia* represents non-specific binding. The results of qRT-PCR were normalized to *Ppia*. For immunoblotting, Gapdh, α-Tubulin and β-Actin are loading controls. Figure C and D shared same loading controls as Figure 1H and 1I. Quantification of protein expression was done by ImageJ (National Institutes of Health, Bethesda, Maryland, USA). *P*-values were calculated by unpaired Student’s t test. Bars represent the mean ± SD. *, *P*<0.05; **, *P*<0.01; ***, *P*<0.001.

Next, we studied whether Stat1 overexpression is sufficient to upregulate Ecm1. Therefore, we transfected AML12 cells with eGFP-Stat1 and eGFP plasmids, respectively. Western blotting revealed that overexpression of Stat1 increased total Stat1 expression, Stat1 phosphorylation at position S727, and Ecm1 expression **(Figure 2E)**. These data consistently demonstrate that Stat1 is required for EGF-EGFR-mediated signal transduction towards Ecm1 expression in hepatocytes.

### EGF induces Stat1 binding to the *Ecm1* promoter for its transcriptional activation

We predicted the binding of Stat1 to the *Ecm1* promoter using JASPAR (https://jaspar.genereg.net/), which showed a putative binding motif ATGGCAGGAAA at −59 ∼ −49bp to the *Ecm1* TSS **(Figure 2F)**. To investigate the regulatory role of Stat1 in *Ecm1* transcription, we constructed a luciferase reporter plasmid containing the corresponding *Ecm1* promoter region (−339 ∼ +161bp) and transfected it into HEK293T cells. Overexpression of Stat1 remarkably increased luciferase activity of the reporter construct **(Figure 2G)**, suggesting that Stat1 binding to the *Ecm1* promoter directly regulates gene transcription. Next, we functionally validated the binding of Stat1 to the predicted binding motif located in the *Ecm1* gene promoter region between −155bp and +120bp in MPHs by chromatin immunoprecipitation (ChIP) qRT-PCR experiments, and the constitutive binding activity was significantly increased by EGF treatment **(Figure 2H)**. The ChIP PCR-amplified products were visualized on 2% agarose gel **(Figure 2I).** Similar results were also obtained in AML12 cells **(Suppl. Fig. 7B, C)**.

These results indicate that EGF induces binding of Stat1 to the proximal promoter region (typically within 250bp upstream of the TSS) of the *Ecm1* gene, thereby activating its transcription.

### IFNγ inhibits EGF/EGFR-mediated Ecm1 expression

IFNγ is a strong modulator of Stat1 signaling (Platanias, 2005) and is found to inhibit HSC activation (Weng *et al*, 2007) but induce hepatocyte apoptosis and hepatic inflammation in CLD (Horras *et al*, 2011). This prompts us to investigate how IFNγ regulates Ecm1 expression in hepatocytes. Treatment of MPHs with IFNγ decreased Ecm1 expression on mRNA and protein levels **(Figure 3A)**. Moreover, IFNγ abrogated EGF-induced Egfr activation and Ecm1 expression, as determined by qRT-PCR and immunoblotting analysis **(Figure 3A)**. EGF and IFNγ can activate Stat1 pathway, but the outcome in the hepatocytes is distinct between them. Although EGF and IFNγ both phosphorylate Stat1 at S727, the difference is that IFNγ requires phosphorylation at Y701 prior to the S727 phosphorylation event (Sadzak *et al*, 2008), whereas EGF is able to directly phosphorylate Stat1 at S727 via EGFR (Zhang *et al*, 2004). In detail, immunoblot analysis of lysates from MPHs treated with EGF or IFNγ for different time intervals displayed that IFNγ induced phosphorylation of Stat1 at Y701 within 5min, which was followed by phosphorylation of S727 at between 10 to 30min **(Figure 3B)**. In contrast, EGF-mediated S727 phosphorylation of Stat1 arose rapidly, whereas Y701 phosphorylation did not take place **(Figure 3B)**. Furthermore, EGF transiently induced Egfr phosphorylation at Y1068 within 5min, therewith initiating its downstream signaling towards Stat1. Moreover, IFNγ inhibits total Egfr expression and was not able to activate Egfr **(Figure 3B)**. ChIP qRT-PCR experiments indicated that IFNγ treatment did not increase Stat1 binding to the *Ecm1* gene promoter in MPHs **(Figure 3C)**. This suggests that unlike Stat1 phosphorylated at S727 alone, Stat1 phosphorylated at both Y701 and S727 cannot bind to the *Ecm1* gene promoter to induce its transcription. Moreover, the addition of IFNγ significantly reduced EGF-induced binding of Stat1 to the *Ecm1* gene promoter as evidenced by the fold enrichment in the ChIP assay **(Figure 3D)**. Next, we administered IFNγ to mice via i.p. injection (400μg/kg BW, for 4 days). Expression of Ecm1 and Egfr were significantly reduced in the IFNγ-treated group compared to controls, as determined by qRT-PCR and immunoblotting **(Figure 3E)**.

**Figure 3.**
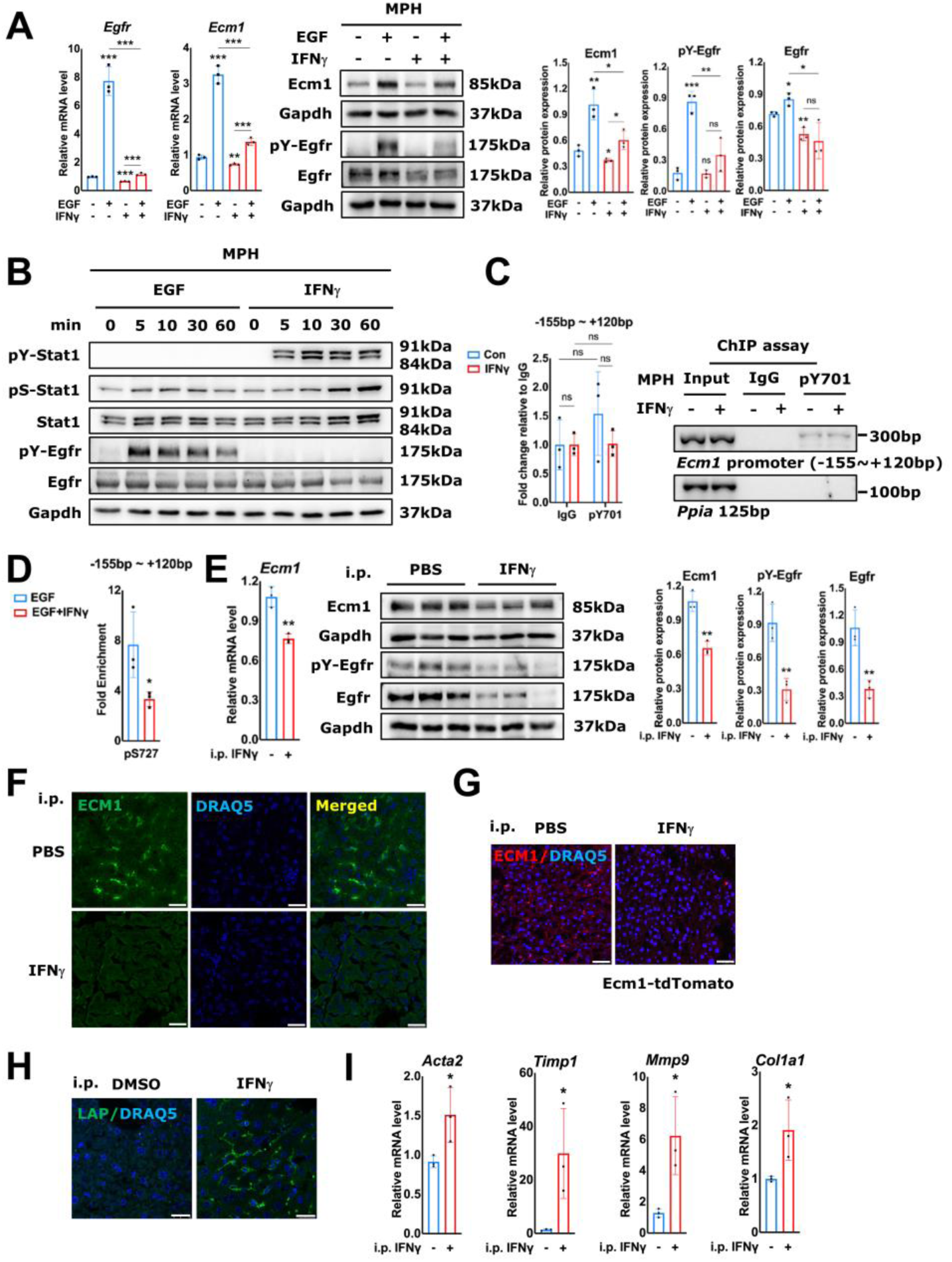
IFNγ abrogates EGF-EGFR-maintained Ecm1 expression. **(A)** qRT-PCR and immunoblotting showing the impact of IFNγ on Ecm1 and Egfr expression in EGF-treated MPHs. **(B)** Immunoblotting showing the effect of EGF or IFNγ on target proteins in MPHs at different time points. **(C)** ChIP qRT-PCR displaying the effect of IFNγ treatment on the binding of p-Stat1 Y701 to the *Ecm1* gene promoter in MPHs. ChIP PCR amplified products were separated on 2% agarose gel. The fragment positions “-155bp ∼ +120bp’’ are calculated relative to the *Ecm1* gene TSS. Rabbit IgG-bound chromatin served as negative control. *Ppia* represents non-specific binding. **(D)** Fold enrichment from ChIP qRT-PCR showing the effect of IFNγ treatment on EGF-induced Stat1 binding to the *Ecm1* gene promoter in MPHs. **(E)** qRT-PCR and immunoblotting showing target genes or proteins in liver tissue from PBS or IFNγ-treated mice. PBS was administered as placebo. **(F)** Representative IF staining for Ecm1 expression in the liver tissue from PBS or IFNγ-treated mice. DRAQ5 fluorescent probe stains DNA. **(G)** Representative tdTomato fluorescence of liver tissue from Ecm1-tdTomato mice treated with PBS or IFNγ. **(H)** Representative IF staining for LAP-D R58 in liver tissues from PBS or IFNγ-treated mice. **(I)** qRT-PCR showing the impact of IFNγ on *Acta2*, *Timp1*, *Mmp9*, *Col1a1* expression in liver tissues from IFNγ-treated mice. Scale bar, 25µm. The results of qRT-PCR are normalized to *Ppia*. Gapdh was used as loading control. Quantification of protein expression was done by ImageJ (National Institutes of Health, Bethesda, Maryland, USA). *P*-values were calculated by unpaired Student’s t test. Bars represent the mean ± SD. *, *P*<0.05; **, *P*<0.01; ***, *P*<0.001.

The decrease in Ecm1 expression was further confirmed by IF staining **(Figure 3F)**. Additionally, Ecm1 was markedly reduced in Ecm1-tdTomato transgenic mice, as observed by tdTomato fluorescence under the same IFNγ treatment conditions **(Figure 3G)**. In line with our hypothesis, IFNγ injection also enhanced LAP-D R58 staining in the liver tissue (**Figure 3H**), indicating upregulated LTGF-β1 activation. Finally, increased expression of fibrotic markers *Acta2*, *Timp1*, *Mmp9*, and *Col1a1* was present in livers of IFNγ-injected mice (**Figure 3I**).

In conclusion, these results imply that IFNγ disrupts Ecm1 homeostasis maintained by EGF-EGFR signaling through inhibiting Egfr expression, and blocks binding of p-Stat1 S727 to the *Ecm1* gene promoter by phosphorylating Stat1 at Y701.

### IFNγ inhibits Ecm1 expression through Nrf2 activation

In CLD, IFNγ promotes inflammation and induces ROS (Lee *et al*, 2007; Miyawaki *et al*, 2019), which leads to Nrf2 activation (Jaeschke & Woolbright, 2012). The *in silico* analysis predicted Nrf2 binding sites at the *Ecm1* gene promoter, indicating Nrf2 may negatively regulate *Ecm1* transcription. Consistent with current knowledge, IFNγ treatment induced ROS-related gene expression, such as *Nos2*, *Cybb*, *Nrf2*, *Nqo1*, *Hmox1*, and *Nox4*, both in cultured cells and in mice **(Figure 4A-B)**. Upregulated ROS gene expression also indicates increased ROS activity, as measured by the fluorescent H2DCFDA, which is converted by intracellular oxidation (488/525 nm) in MPHs treated with IFNγ **(Figure 4C)**. Finally, IF staining of tissue sections demonstrated nuclear accumulation of Nrf2 in the livers of IFNγ-treated mice **(Figure 4D),** however, no differences for total Nrf2 protein expression were evident based on immunoblots **(Suppl. Fig. 9A)**. Next, we treated MPHs with EGF and/or IFNγ and prepared nuclear and cytoplasmic lysates for immunoblot analysis. It shows that the Nrf2 signal in the nuclear fraction was strongly induced upon IFNγ treatment **(Figure 4E)**. IFNγ dependent Nrf2 nuclear accumulation in MPHs was additionally confirmed by IF staining for Nrf2 **(Figure 4F)**. To further investigate whether IFNγ-mediated Ecm1 downregulation is dependent on Nrf2, we depleted Nrf2 with RNAi in the setting of IFNγ treatment in MPHs **(Suppl. Fig. 9B)**. The data shows that Nrf2 knockdown rescued IFNγ-mediated Ecm1 downregulation **(Figure 4G)**.

**Figure 4.**
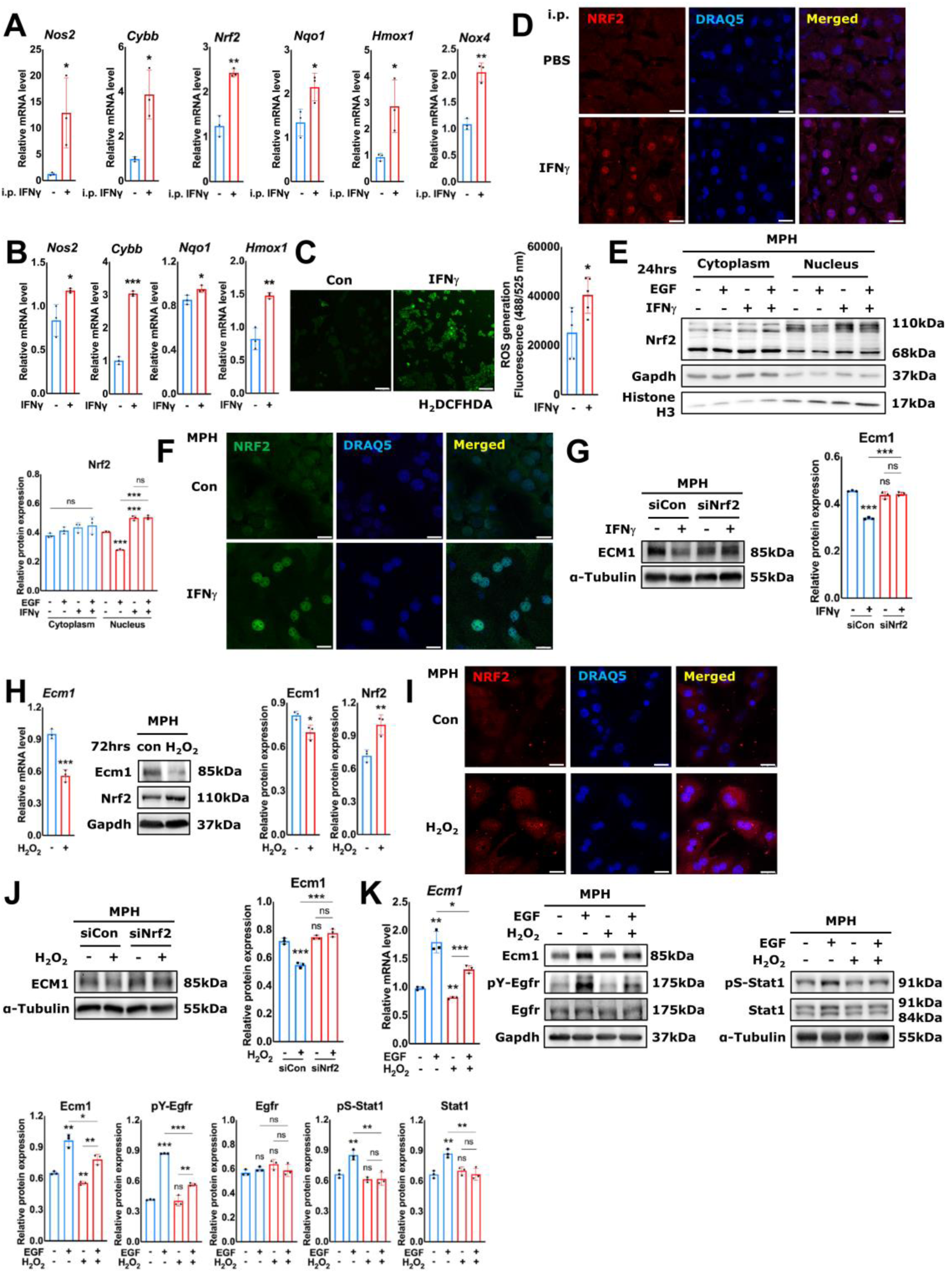
IFNγ inhibits Ecm1 expression through Nrf2. **(A)** qRT-PCR for mRNA expression levels of *Nos2*, *Cybb*, *Nrf2*, *Nqo1*, *Hmox1* and *Nox4* in liver tissues from IFNγ-treated mice. **(B)** qRT-PCR for mRNA expression levels of *Nos2*, *Cybb*, *Nqo1*, *Hmox1* in IFNγ-treated MPHs. **(C)** ROS generation fluorescence detection in the MPHs treated with IFNγ. **(D)** Representative IF staining for Nrf2 expression in liver tissues from PBS or IFNγ-treated mice. Scale bar, 12.5µm. **(E)** Immunoblotting for cytoplasmic and nuclear localization of Nrf2 in MPHs treated with EGF and/or IFNγ. **(F)** Representative IF staining showing expression and localization of Nrf2 in IFNγ-treated MPHs. Scale bar, 25µm. **(G)** Immunoblotting showing the impact of *Nrf2* knockdown on Ecm1 expression in IFNγ-treated MPHs. **(H)** qRT-PCR and immunoblotting showing the impact of H_2_O_2_ treatment on Ecm1 and Nrf2 expression in MPHs. **(I)** Representative IF staining showing Nrf2 expression in H_2_O_2_-treated MPHs. Scale bar, 25µm. **(J)** Immunoblotting showing the impact of knockdown *Nrf2* on Ecm1 expression in H_2_O_2_-treated MPHs. **(K)** qRT-PCR and immunoblotting showing the effect of H_2_O_2_ on target genes or proteins in EGF-treated MPHs. The results of qRT-PCR are normalized to *Ppia*. Gapdh/α-Tubulin and Histone H3 are loading controls for cytoplasmic and nuclear proteins, respectively. DRAQ5 fluorescent probe stains DNA. Quantification of protein expression was done by ImageJ (National Institutes of Health, Bethesda, Maryland, USA). *P*-values were calculated by unpaired Student’s t test. Bars represent the mean ± SD. *, *P*<0.05; **, *P*<0.01; ***, *P*<0.001.

Given that Nrf2 is activated by ROS (Jaeschke & Woolbright, 2012), we treated MPHs with hydrogen peroxide (H_2_O_2_) to directly induce oxidative stress. With this experiment, we demonstrated an alternative route of ROS-mediated Nrf2 activation with the same effect on Ecm1 mRNA and protein expression downregulation **(Figure 4H, I)**. Depleting Nrf2 in this setting (**Suppl. Fig. 9C)** rescued the H_2_O_2_-mediated Ecm1 downregulation **(Figure 4J)**. In addition, H_2_O_2_ inhibited EGF-induced Y1068 phosphorylation of the Egfr, S727 phosphorylation of Stat1, total Stat1-, as well as Ecm1 expression **(Figure 4K)**. Bright field photos of MPH treated with H_2_O_2_ indicate that the cells were in good conditions during the experiment **(Suppl. Fig. 9D)**, excluding the assumption that the observed effects are relying on cellular damage.

These results together suggest that IFNγ inhibits Ecm1 expression in hepatocytes through Nrf2 activation. Given its potential to bind the *Ecm1* promoter, Nrf2 is a candidate regulator for *Ecm1* transcription modulation.

### Nrf2 inhibits Ecm1 expression through negative regulatory binding to its promoter

To further confirm the inhibitory effect of Nrf2 on Ecm1 expression, we used a specific agonist of Nrf2, Oltipraz (OPZ). The data showed that OPZ, as expected, promoted Nrf2 expression on mRNA and protein levels **(Figure 5A)** and induced its nuclear accumulation in MPHs **(Figure 5B)**. Ecm1 expression was significantly downregulated upon OPZ treatment in MPHs **(Figure 5C)**. Importantly, knockdown of Nrf2 effectively mitigated the OPZ-induced reduction on Ecm1 levels **(Figure 5D),** suggesting that despite several other known functions of OPZ, the observed effects are specifically relying on stabilizing Nrf2. Further, the OPZ treatment effects could be phenocopied by Nrf2 overexpression, which as well prevented EGF-induced Ecm1 expression **(Figure 5E)**. Finally, OPZ-mediated Nrf2 activation was able to abrogate EGF-driven upregulation of Ecm1 expression **(Figure 5F)**.

**Figure 5.**
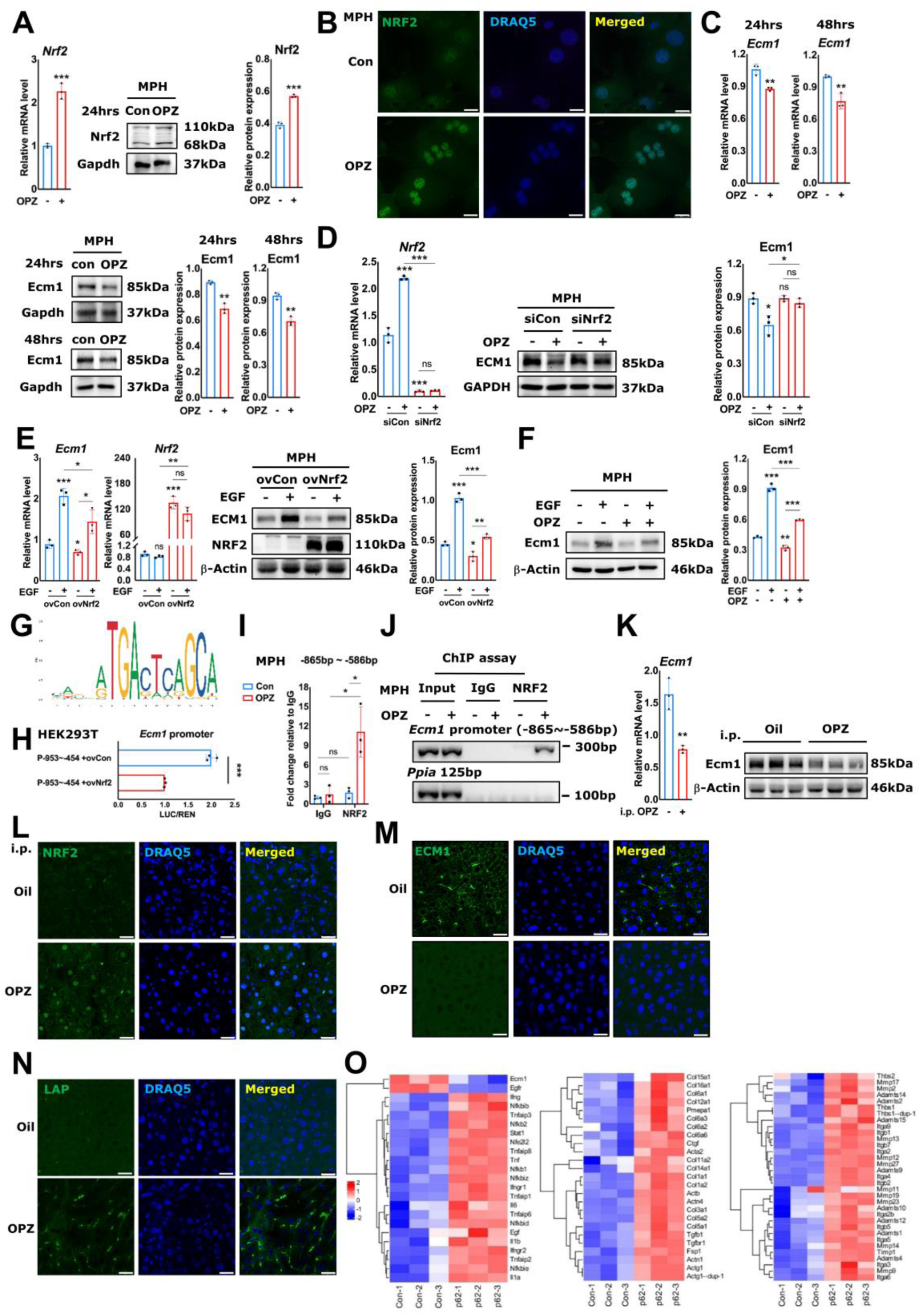
Nrf2 inhibits Ecm1 expression through negative regulatory binding to its promoter. **(A)** qRT-PCR and immunoblotting showing the effect of OPZ treatment on Nrf2 expression in MPHs. **(B)** Representative immunofluorescence staining for Nrf2 expression in OPZ-treated MPHs. Scale bar, 25µm. **(C)** qRT-PCR and immunoblotting displaying the effect of OPZ treatment on Ecm1 expression in MPHs at different time points. **(D)** qRT-PCR and immunoblotting showing the effect of Nrf2 knockdown on Ecm1 expression in OPZ-treated MPHs. **(E)** qRT-PCR and immunoblotting showing the effect of Nrf2 overexpression on Ecm1 and Nrf2 expression in EGF-treated MPHs. **(F)** Immunoblotting displaying the effect of OPZ treatment on Ecm1 expression in EGF-treated MPHs. **(G)** Predicted binding motif of Nrf2 to the *Ecm1* promoter by Jaspar. **(H)** Luciferase reporter assay analyzing the activity of *Ecm1* promoter (−953 ∼-454bp) in HEK293T cells transfected with or without Nrf2 plasmid. **(I)** ChIP qRT-PCR showing the effect of OPZ treatment on binding of Nrf2 to the *Ecm1* gene promoter in MPHs. The fragment positions “-865bp ∼ −586bp’’ were calculated relative to the *Ecm1* gene TSS. Rabbit IgG-bound chromatin served as negative control. **(J)** The amplified products of ChIP PCR were separated on 2% agarose gel. *Ppia* represents non-specific binding. **(K)** qRT-PCR and immunoblotting showing the expression of Ecm1 in OPZ-treated mice. Olive oil was administered as placebo. **(L-N)** Representative IF staining showing Nrf2, ECM1 and LAP-D R58 expression in liver tissues from OPZ-treated mice. **(O)** mRNA expression of target genes in liver tissue from p62-KO mice injected with/without adenovirus p62, extracted from the GEO Dataset GSE134188. The results of qRT-PCR are normalized to *Ppia*. Gapdh and β-Actin are loading controls. DRAQ5 fluorescent probe stains DNA. Scale bar, 25µm. Quantification of protein expression was done by ImageJ (National Institutes of Health, Bethesda, Maryland, USA). *P*-values were calculated by unpaired Student’s t test. Bars represent the mean ± SD. *, *P*<0.05; **, *P*<0.01; ***, *P*<0.001.

To investigate whether Nrf2 prevented *Ecm1* expression through its transcription inhibition, we constructed a luciferase reporter plasmid comprising *Ecm1* promoter (−953 ∼ −454bp), which contained the predicted Nrf2 binding site at positions −608 ∼ - 594bp of the *Ecm1* gene promoter (**Figure 5G**, predicted binding motif antisense GGACATGACTCAGAA) and was transfected into HEK293T cells. Overexpression of Nrf2 markedly reduced the luciferase activity of the reporter construct **(Figure 5H)**, indicating that Nrf2 directly and negatively regulates the *Ecm1* promoter. We next performed ChIP qRT-PCR for the region −865bp to −586bp of *Ecm1* gene promoter, in OPZ-treated MPHs, and functionally proved that OPZ-induced Nrf2 binds directly to the *Ecm1* gene promoter **(Figure 5I)**. Amplification products were visualized on 2% agarose gel **(Figure 5J)**.

To validate the effect of Nrf2 on Ecm1 expression *in vivo*, we performed i.p. injection of OPZ (150mg/kg BW) into WT mice once per week for 2 weeks. It showed that OPZ treatment resulted in Ecm1 downregulation at the mRNA and protein levels **(Figure 5K)**. IF staining confirmed the nuclear translocation of Nrf2, reduction of Ecm1 expression, and LTGF-β activation in OPZ-treated mice **(Figure 5L-N)**. Next, we extracted mRNA expression data from GEO dataset GSE134188 (He *et al*, 2020) of the target genes involved in the Ecm1 expression regulation network from a mouse model of hyperactivated Nrf2. Specifically, liver-specific autophagy adaptor p62/Sqstm1 (p62)-KO mice were injected with/without adenovirus p62 and were analyzed 7 days after injection. In these mice, p62 expression sequesters inhibitory Kelch-like ECH-associated protein 1 (KEAP1) from Nrf2, which results in activation and subsequent nuclear accumulation of activated Nrf2 (He *et al*., 2020). The analysis demonstrates that the pro-inflammatory signals (*Ifng*, *Ifngr*, *Tnf*, *Nfkb*, *Il1*, *Il6*, et.al), *Stat1*, and *Nrf2* (*Nfe2l2*) were notably increased, while *Egfr* and *Ecm1* were decreased **(Figure 5O, left penal)**. In consequence, TGF-β signaling was hyperactivated, here documented by upregulated expression of *Tgfbr1*, *Ctgf*, and *Pmepa1*, which resulted in HSC activation (upregulated *Acta2*), and fibrogenesis, as evident from upregulated expression of fibrogenic genes (*Col1a1*, *Col1a2*, *Col3a1, Fsp1*) **(Figure 5O, middle penal)** and genes related to extracellular matrix remodelling (*Timp1*, *Integrins*, *Adamts*, *Thbs*, *Mmps*) **(Figure 5O, right penal)**.

### Activation of IFNγ/NRF2 signaling and downregulation of EGFR/ECM1 in various mouse models of hepatic injury

Given the considerable downregulation of Ecm1 in several mouse models with progressive liver fibrosis (Fan *et al*., 2019), we further investigated the regulatory network of hepatic Ecm1 expression by evaluating mRNA expression of target genes from the GEO dataset. Mice were challenged with CCl_4_ (twice per week for 6 weeks, GSE222576) (Hammad *et al*, 2023) **(Figure 6A)**, underwent surgical BDL (7 days after ligation, GSE166867) (Holland *et al*, 2022) **(Figure 6B)**, knocked out of the *Fxr* gene, which encodes a nuclear receptor regulating bile acid and cholesterol homeostasis (GSE76163) (Ijssennagger *et al*, 2016) **(Figure 6C)**, given 0.1% DDC diet (for 1 week, GSE193850) (Li *et al*., 2023) **(Figure 6D)**, or fed with CD-HFD (for 3 months, GSE232182) **(Figure 6E)**. Bulk RNA-seq and scRNA-seq analysis illustrated the downregulation of *Ecm1*, *Egfr*, along with the upregulation of *Ifngr1*, *Ifngr2*, *Nfe2l2* and its target genes, including antioxidant enzymes cytochrome P450 family 1 subfamily B member 1 (*Cyp1b1*) and NAD(P)H:quinone oxidoreductase 1 (*Nqo1*). Additionally, TGF-β signaling was markedly activated in these mouse models, as evidenced by the enhanced expression of relevant genes *Tgfbr1*, *Ctgf*, and *Pmepa1*, leading to HSC activation (upregulated *Acta2*), and fibrosis progression (upregulated *Fsp1*, *Actn1*, *Col1a1*, *Col1a2*, *Col3a1*) **(Figure 6A-E)**. Moreover, immunoblotting and IF/IHC staining further confirmed suppression of EGFR activation, downregulation of Ecm1 expression, and activation of Nrf2 signals in liver tissue of mice treated for 6-weeks with CCl_4_ **(Figure 6F, G; Suppl. Fig. 10)** or fed a Western diet (WD) for 12-weeks **(Figure 6H, I)**, both supporting our findings in CLD models.

**Figure 6.**
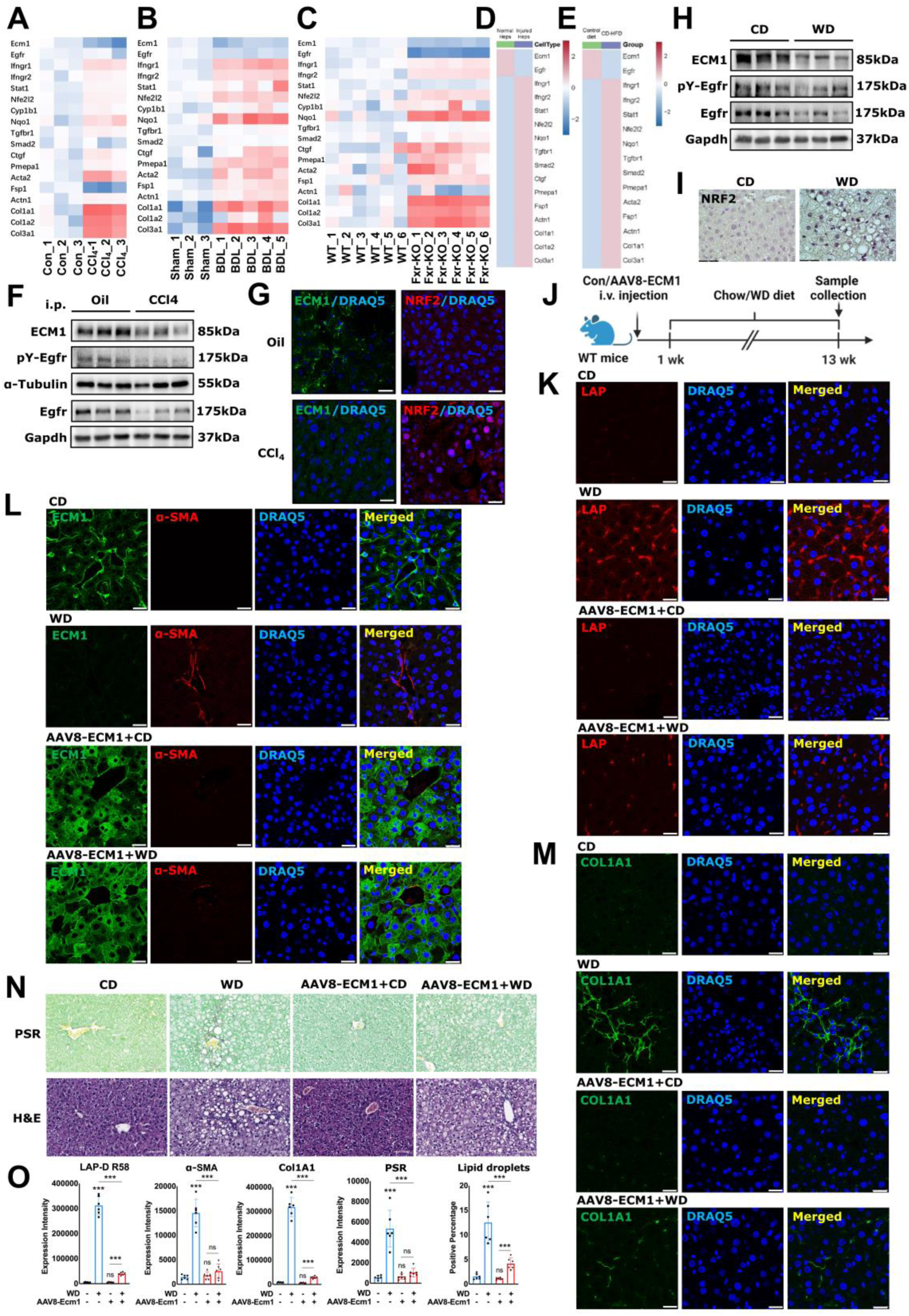
Activation of IFNγ/NRF2 signaling and downregulation of EGFR/ECM1 in various mouse models of hepatic injury. **(A-E)** Heatmaps of relative mRNA expression for target genes in CCl_4_-injured (twice per week for 6 weeks), BDL surgical (7 days after ligation), *Fxr*-KO, DDC diet (for 1 week) and CD-HFD fed (for 3 months) mouse models, extracted from the GEO Dataset GSE222576, GSE166867, GSE76163, GSE193850 and GSE232182, respectively. **(F)** Immunoblotting for Ecm1, pY-Egfr and Egfr in liver tissues from oil or CCl_4_-treated mice. **(G)** Representative IF staining for Ecm1 and Nrf2 in liver tissues from oil or CCl_4_-treated mice. **(H)** Immunoblotting for Ecm1, pY-Egfr and Egfr in liver tissues from CD or WD-fed mice. **(I)** Representative IHC staining for Nrf2 in liver tissues from CD or WD-fed mice. **(J)** Schematic illustration of *in vivo* experiment with AAV8-ECM1 (i.v.) and WD-fed WT mice. **(K-M)** Representative IF staining for LAP-D R58, Ecm1, a-SMA and Col1A1 in liver tissues from CD or WD-fed mice treated with control or AAV8-ECM1. DRAQ5 fluorescent probe stains DNA. Scale bar, 25µm. **(N)** Representative H&E and PSR staining for liver tissues from CD or WD-fed mice treated with control or AAV8-ECM1. Scale bar, 70µm. **(O)** Quantification of staining for LAP-D R58, a-SMA, Col1A1 and PSR, and for lipid droplets according to the white bubbles in the H&E staining. Gapdh, α-Tubulin are loading controls. *P*-values were calculated by unpaired Student’s t test. Bars represent the mean ± SD. *, *P*<0.05; **, *P*<0.01; ***, *P*<0.001.

### Overexpression of ECM1 inhibits WD-induced liver injuries

MASLD is the most common liver disease, with its severe form, MASH, characterized by liver inflammation and fibrosis, which can progress to cirrhosis and HCC (Wang *et al*, 2024). Lipid droplets that accumulate during the progression of MASLD change the composition of free fatty acids, triglycerides, cholesterol, and phospholipids, thereby inducing oxidative stress in the liver (Wang *et al*., 2024). Since we found the activation of IFNγ/ROS signaling and downregulation of EGFR/ECM1 in mouse models fed with HFD/WD, we investigated whether ECM1 overexpression prevents WD-induced liver fibrosis. Therefore, we treated 8-week-old WT mice with control or AAV8-ECM1 for 7 days prior to 12 weeks of WD feeding (Chow diet (CD) was used as control) **(Figure 6J)**. WD treatment resulted in significant activation of latent TGF-β1 (LTGF-β1), as evidenced by the increased expression of LAP-D R58, a breakdown product of the TGF-β1 LAP deposited in the ECM upon LTGF-β1 activation (**Figure 6K**). Additionally, WD-fed mice showed upregulation of HSC-specific fibrosis markers, including α-SMA and Col1a1 (**Figures 6L, M**), and exhibited moderate hepatic fibrosis and liver injury, confirmed by PSR and H&E staining (**Figure 6N**). Remarkably, all these pathological changes were significantly attenuated by AAV8-ECM1 treatment (**Figures 6K-O**). Successful ECM1 overexpression was confirmed via IF staining (**Figure 6L**). Interestingly, AAV8-ECM1 also appeared to reduce fat accumulation in the liver tissue of WD-treated mice, as indicated by the lipid droplets of the white bubbles in the H&E staining (**Figure 6N, O**). However, whether this effect is mediated through TGF-β signaling or alternative pathways warrants further investigation. Taken together, ECM1 overexpression effectively prevents liver fibrosis and injuries induced by WD treatment, suggesting its potential benefit in MASLD progression.

### Inhibited EGFR, activated IFN**γ/**NRF2, and decreased ECM1 expression aggravate the progression of fibrotic liver diseases

To further verify the relevance of reduced ECM1 expression and its link to inhibited EGF signaling, activated NRF2, and enhanced TGF-β signaling in CLD patients, we performed IF or IHC staining for ECM1, EGFR, NRF2, and TGF-β1 LAP-D R58 in the liver tissue from patients with early stages of liver fibrosis (F1-F2 fibrosis) or more severe disease stages of F3-F4 fibrosis. The expression of ECM1 and EGFR decreased at the F3-F4 stage compared to F1-F2 patients, whereas nuclear NRF2 and latent TGF-β activation (LAP-D R58) increased considerably with disease progression **(Figure 7A)**. Quantification of the stainings implicated a positive correlation between the expression of EGFR and ECM1 (correlation coefficient 0.9710) **(Figure 7B),** whereas there is a negative correlation between NRF2 and ECM1 (correlation coefficient - 0.835) **(Figure 7C)**. This was further confirmed by analysing the GEO dataset GSE49541 (Moylan *et al*, 2014; Murphy *et al*, 2013) that comprises mRNA expression data of liver tissue from MASLD patients with mild (n=40) or advanced (n=32) fibrosis. Gene set variation analyses (GSVA) revealed significant activation and upregulation of IFNγ-, NRF2-, and TGF-β-related pathways in patients with advanced liver fibrosis, including, among others, the oxidative stress pathway, the TGF-β signaling pathway, the inflammatory response, type II interferon signaling, and the NRF2 pathway **(Figure 7D)**. We further analyzed the expression of key genes involved in these pathways, which showed that the mRNA expression of *IFNGR1*, *STAT1,* and *NFE2L2* target genes, including *CYP1B1* and *NQO1*, did increase with disease severity, while mRNA levels of *ECM1* were decreased, leading to enhanced TGF-β signaling (*TGFBR1, SMAD2, CTGF, PMEPA1*) (Link *et al*., 2024), supported by the presence of upregulated HSC activation marker *ACTA2* (encoding α-SMA) and fibrogenic gene expression, including collagens, such as *FSP1*, *ACTN1*, *COL1A1*, *COL1A2*, and *COL3A1* in the patients with more advanced fibrosis **(Figure 7E)**. Moreover, scRNA-seq dataset GSE174748 including two normal liver samples and two MASLD cirrhotic liver samples was analyzed (Filliol *et al*., 2022). Unsupervised clustering classified cells into 14 different clusters **(Suppl. Fig. 11A)**. Based on the marker genes illustrated in **Suppl. Fig. 11B**, we were able to classify the cell clusters into various categories including B cell, Macrophage, T cell, Choloangiocyte, Endothelial cell, HSC and Hepatocyte **(Figure 7F)**. Differential expression analysis in the hepatocytes between healthy and cirrhotic livers was performed **(Figure 7G)**. Heatmap showed that in the hepatocytes of cirrhotic livers, the expression of *ECM1*, *EGFR* were significantly deseased whereas *IFNGR1* and the downstream targets of NRF2, *CYP1B1* and *NQO1*, were dramatically increased. Additionally, the enhancement of TGF-β signaling was evidenced by upregulated expression of *TGFBR1* in the hepatocytes from cirrhotic livers **(Figure 7H)**. Similar results were also confirmed in MASLD/MASH patients from the aforementioned dataset GSE202379 (Gribben *et al*., 2024), where downregulation of *ECM1* was accompanied by upregulation of *IFNGR1*, *NFE2L2* and *TGFBR1*, as well as *COL1A1*, compared to healthy donors **(Figure 7I)**.

**Figure 7.**
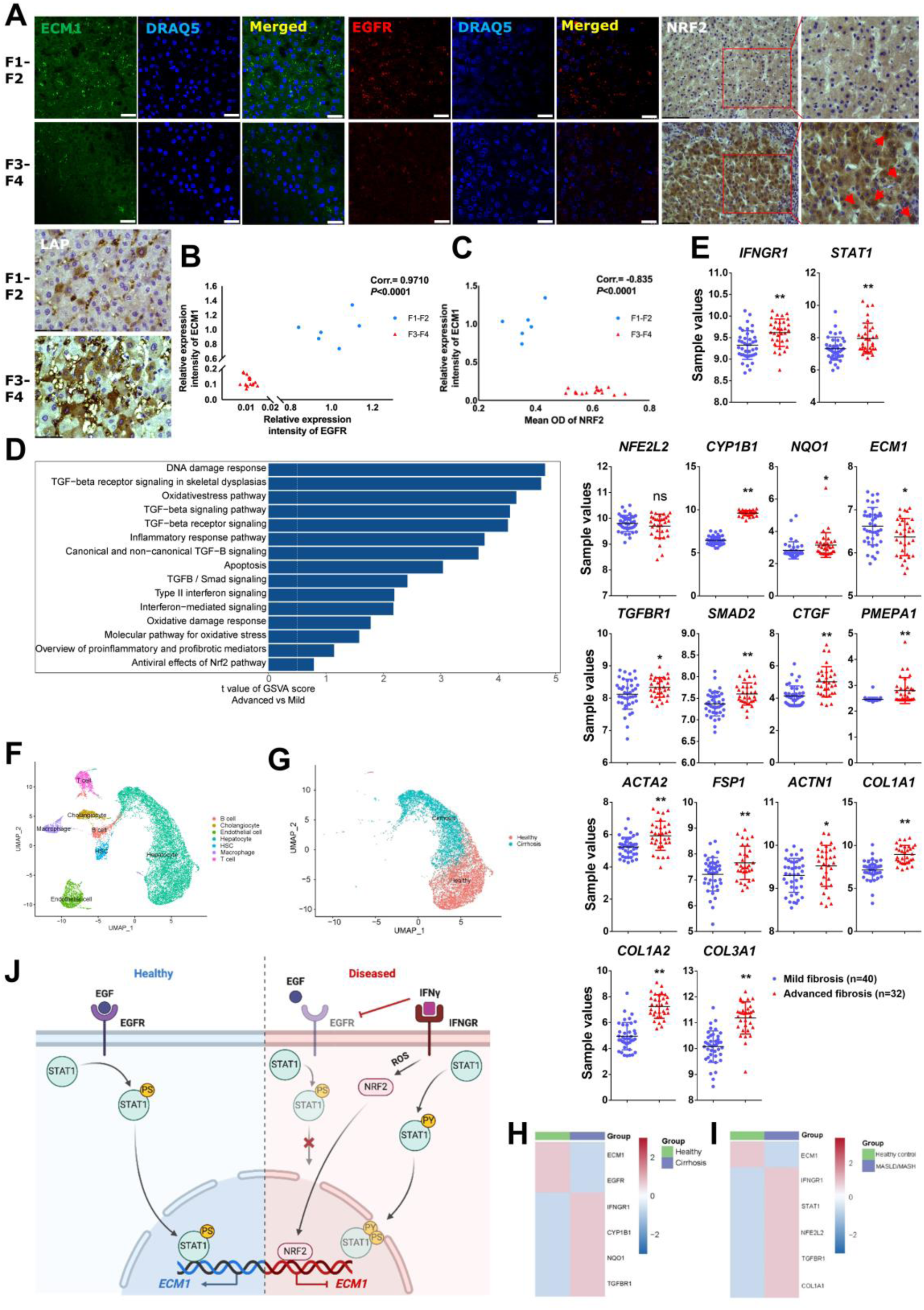
Activated IFN**γ**/NRF2 and decreased ECM1 expression aggravate the progression of fibrotic liver diseases. **(A)** Representative IF or IHC staining for ECM1, EGFR, NRF2, and TGF-β1 LAP-D R58 in F1-F2 fibrosis and F3-F4 fibrosis patients. DRAQ5 fluorescent probe stains DNA. Scale bar, IF 25µm, IHC 87µm. **(B-C)** Correlation analysis of EGFR/NRF2 and ECM1 expression levels in the liver tissue of patients with F1-F2 or F3-F4 fibrosis. **(D)** Differences in pathway activity were evaluated using GSVA scores for each patient with advanced (n = 32) and mild liver fibrosis (n = 40). The analysis reports t-values derived from linear models. **(E)** mRNA expression of target genes in liver tissue from patients with F0-F1 or F3-F4 fibrosis, extracted from GEO Dataset GSE49541. **(F, G)** UMAP visualization of cell clusters. **(H)** Analysis of target genes in the hepatocytes from human normal liver (n=2) and MASLD cirrhotic liver (n=2), extracted from scRNA-seq dataset GSE174748. **(I)** Heatmap of target genes expression in MASLD/MASH patients (GSE202379). **(J)** Scheme depicting the regulation of ECM1 expression in hepatocytes in healthy and diseased liver (Figure created with BioRender.com). *P*-values were calculated by unpaired Student’s t test. Bars represent the mean ± SD. *, *P*<0.05; **, *P*<0.01; ***, *P*<0.001.

In conclusion, (1) in healthy hepatocytes, the EGF/EGFR/STAT1 signaling pathway maintains ECM1 expression. However, (2) upon liver injury, IFNγ accumulates in the liver and abrogates the expression of ECM1 through blocking EGFR expression and (3) promoting NRF2 nuclear translocation, which binds to and negatively regulates the *ECM1* gene promoter **(Figure 7J)**, leading to ECM1 downregulation, LTGF-β activation, and CLD progression. The regulatory mechanism of ECM1 is further validated in various CLD mouse models treated with CCl_4_, BDL, *Fxr*-KO, DDC diet, or CD-HFD. Moreover, ECM1 overexpression effectively attenuates MASLD progression induced by WD treatment.

## Discussion

This study investigated the regulation of ECM1 expression in hepatocytes under various pathophysiological conditions associated with liver disease. Given hepatocytes are the most vulnerable cells in the liver (Gribben *et al*., 2024), ECM1 expression is rapidly and significantly decreased in hepatocytes following injury. Although HSCs also secrete ECM1, which serves as an important marker for their quiescence, the reduction of ECM1 by activated HSCs occurs subsequent to hepatocyte injury (Fan *et al*., 2019; Link *et al*., 2024). This suggests that the downregulation of ECM1 in injured hepatocytes during the early stages of CLD may be a key factor in driving fibrosis progression. Nevertheless, we examined the effects of EGF, IFNγ, and OPZ on *ECM1* expression in human HSC LX-2 cells. However, these factors did not significantly affect *ECM1* expression, indicating that the regulation of ECM1 in HSCs differs from that in hepatocytes and warrants further investigation.

Our results showed that EGF promotes, whereas IFNγ inhibits ECM1 expression. Why is the regulation of ECM1 expression so different despite the fact they both exploit STAT1 signaling? Based on three lines of evidence, we elucidated that IFNγ and EGF-induced STAT1 Ser727 phosphorylation are mechanistically independent events: (1) Tyr701 phosphorylation of STAT1 is necessary for IFNγ-induced STAT1 Ser727 phosphorylation (Sadzak *et al*., 2008). However, Ser727 phosphorylation induced by the p38 mitogen-activated protein kinase (MAPK) is independent of Tyr701 phosphorylation (Kovarik *et al*, 1999). (2) EGF induced Ser727 phosphorylation alone without Tyr701 phosphorylation (Zhang *et al*., 2004), which is consistent with our results **(Figure 3B)**. (3) EGF treatment enhanced STAT1 binding to the *Ecm1* promoter **(Figure 2H, I; Suppl. Fig. 7B, C)**, whereas IFNγ treatment impeded the EGF-promoted binding **(Figure 3C, D)**. It is important to recognize the multipotent effects of STAT1 signaling, which may be regulated by EGF and involve essential physiological functions beyond IFNγ. In mammary carcinoma cells, EGF/STAT1 signaling can antagonize the growth-inhibitory effects of TGF-β signaling via EGFR-AKT-mediated phosphorylation of Smad3 at S208 (Huang *et al*, 2018). Conversely, TGF-β signaling can affect the EGF/STAT1 pathway by downregulating EGFR expression, therewith inhibiting STAT1 activation and reducing its DNA binding ability (Baba *et al*, 2022; Huang *et al*., 2018). We now add the knowledge that, in the liver, EGF/STAT1 can inhibit latent TGF-β activation through ECM1 expression upregulation.

IFNγ, mainly produced by activated T cells and natural killer cells, is an anti-viral, pro-inflammatory and anti-tumorigenic cytokine, and has been reported to have anti-fibrotic properties in hepatic fibrosis associated with chronic HBV infections (Weng *et al*, 2005). One trial showed that nine-month IFNγ treatment improves fibrosis scores in patients through antagonizing TGF-β signaling (Weng *et al*., 2005). *In vitro*, IFNγ inhibits the proliferation and activation of HSCs, thereby inhibiting liver fibrosis (Jeong *et al*, 2006; Weng *et al*., 2007). Nonetheless, IFNγ also induces hepatocyte apoptosis (Tagawa *et al*, 1997), hepatic inflammation, and inhibits hepatocyte proliferation, liver regeneration (Sun *et al*, 2006), contributing to liver fibrogenesis. In viral hepatitis, the hepatic lesions can be attributed to T cell-dependent cytotoxicity against virus-infected hepatocytes, where IFNγ, which is typically elevated in patients with chronic viral liver disease (Fukuda *et al*, 1995), plays a crucial role. When IFNγ was administered to HBV transgenic mice that did not develop hepatitis, hepatic lesions with lymphocytes infiltration were observed (Gilles *et al*, 1992). Moreover, in a methionine and choline-deficient high-fat (MCDHF) diet mouse model, IFNγ was even identified as a pro-fibrotic cytokine (Luo *et al*, 2013). NRF2 is activated by inflammatory mediators such as ROS, fatty acids, nitric oxide and prostaglandins (Saha *et al*, 2020), which is considered to provide a cytoprotective effect. However, excessive activation of NRF2 and a consecutive nuclear accumulation has detrimental effects. In a *Keap1*-null mouse model, no newborns survived after three weeks, possibly due to starvation caused by a hyperkeratotic esophagus and cardia (Wakabayashi *et al*, 2003). Additionally, an association between NRF2 and hepatic steatosis was identified, demonstrating that a specific deletion of NRF2 in hepatocytes reduced the expression of HFD-induced PPARγ and lipid accumulation, thus impairing the progression of MASLD (Li *et al*, 2020). Moreover, pancreatic cancer patients with elevated NRF2 levels have a shorter median survival time (Su *et al*, 2022). Our findings suggest that both, NRF2 and IFNγ integrate in damage stressed hepatocytes towards, among others, downregulation of ECM1. Loss of ECM1 is a kind of negative damage-associated molecular pattern (DAMP) reaction, which results in extracellular decrease in ECM1 availability, subsequent LTGF-β activation. Thus, increased amounts of active TGF-β now can activate HSCs and induce fibrogenesis. In terms of anti-fibrotic therapy, while using IFNγ to inhibit HSC activation or NRF2 agonist in CLD patients, their potential adverse effects on hepatocytes should also be considered.

Overall, the current study highlights the roles of EGF and IFNγ in regulating ECM1 expression in hepatocytes under physiological and pathological conditions. For the clinical applicability of IFNγ and NRF2 agonist, the potential hepatotoxicity resulting from IFNγ and NRF2 over-accumulation should be carefully evaluated, as they significantly inhibit ECM1 expression in hepatocytes. ECM1 has the potential to be developed as an anti-fibrotic agent, especially in combination with IFNγ and NRF2 agonist. It will be interesting to further investigate whether ECM1-derived therapies constitute an effective treatment route for CLD.

## Materials and Methods

Human samples, mouse experiments, primers, and reagents used in this study, as well as methodological details can be found in the supplementary materials.

This study includes no data deposited in external repositories.

## Acknowledgements

We acknowledge the support of the LIMa Live Cell Imaging at Microscopy Core Facility Platform Mannheim (CFPM).

## Abbreviations

ADAMTS1: A disintegrin and metalloproteinase with thrombospondin motifs 1
BDL: Bile duct ligation
BW: Body weight
CCl_4_: Carbon tetrachloride
CD: Chow diet
CD-HFD: Choline-deficient high-fat diet
ChIP: Chromatin immunoprecipitation
CLD: Chronic liver disease
CYP1B1: cytochrome P450 family 1 subfamily B member 1
DDC: 3,5-diethoxycarbonyl-1,4-dihydrocollidine
ECM1: Extracellular matrix protein 1
EGF: Epidermal growth factor
EGFR: Epidermal growth factor receptor
GEO: Gene Expression Omnibus
H_2_O_2_: Hydrogen peroxide
HBV: Hepatitis-B-virus
HCC: Hepatocellular carcinoma
HPHs: Human primary hepatocytes
HSCs: Hepatic stellate cells
IF: Immunofluorescence
IFNγ: Interferon gamma
IHC: Immunohistochemistry
i.p.: Intraperitoneal
KEAP1: Kelch-like ECH-associated protein 1
LAP: Latency-associated peptide
LTGF-β: Latent TGF-β
MELD: Model for End-Stage Liver Disease
MMP: Matrix metalloproteinase
MPHs: Mouse primary hepatocytes
MASLD: Metabolic dysfunction-associated steatotic liver disease
MASH: Metabolic dysfunction-associated steatohepatitis
NQO1: NAD(P)H:quinone oxidoreductase 1
NRF2: Nuclear factor erythroid 2-related factor 2
OPZ: Oltipraz
PSR: Picrosirius Red
qRT-PCR: Quantitative real-time PCR
ROS: Reactive oxygen species
scRNA-seq: single-cell RNA sequencing
siRNA: Small interfering RNA
S727: TSP-1 TSS
Serine727: Thrombospondin 1 Transcription start site
UMAP: Uniform manifold approximation and projection
WD: Western diet
Y701: Tyrosine701

## Author contributions

Conceptualization: SD, SW

Methodology: YL, SW, WF, CH, SH, CT, ZCN, YY, PE, WP, KG, RF, HL, CS

Investigation: YL, SW, CH, WF, SH, YY, FL, CT, ZCN, LB, CG, PE, CM, HL, CS

Visualization: YL, SW, CH

Funding acquisition: SD

Project administration: SD, SW, YL

Supervision: SW, SD

Writing - Original: YL, SW, SD

Writing - review: YL, SW, SD, PTD, RL, CH, FL, ZCN, CM, BS, HD, ME, CG, HW

SW and SD acted as guarantors.

