## Supplementary material for "Balance Between EGF/STAT1 and IFNγ/NRF2 Signaling Controls ECM1 Expression and Determines Hepatic Homeostasis versus Chronic Liver Disease"

### Supplementary materials and methods

#### Patients

6 F1-F2 fibrotic liver tissues were collected by biopsy; 16 F3-F4 fibrotic liver tissues were obtained from patients following liver transplantation; 76 paraneoplastic liver tissues were obtained from HCC patients with or without cirrhosis during surgery at the Beijing You'an Hospital, Affiliated with Capital Medical University. The study protocol was approved by local Ethics Committees (Jing-2015-084, and 2017-584N-MA). Written informed consent was obtained from patients or their representatives.

#### Animals

Male C57BL/6J mice (8 to 10 weeks old) were purchased from the Janvier Lab. Ecm1-tdTomato mice were purchased from Transgenic Core Facility, University of Copenhagen using CRISPR-Cas9 targeted integration of a DNA template at the 5' end of the mouse *Ecm1* gene. All animals weighed 22-25g at the beginning of the corresponding experiments and were allowed to acclimatize to controlled conditions of temperature (23 ± 2°C), humidity (35 ± 5%) and a 12hrs light–dark cycle in the animal house at the Universitätsmedizin Mannheim for at least 1 week. They were provided with standard laboratory chow and water ad libitum and housed in laboratory cages. The mice were divided randomly into groups (n=3) and were injected with EGF, erlotinib, IFNγ or OPZ. EGF, erlotinib, IFNγ and OPZ were dissolved in PBS (14190169, ThermoFisher), DMSO (41639-500ML, Sigma-Aldrich), PBS and olive oil, respectively. For EGF injection, mice were given 200μg/kg body weight (BW) EGF through tail vein with one dosage for 24h or two dosages for two consecutive days. For erlotinib injection, mice were administered intraperitoneally with 40mg/kg BW erlotinib or DMSO, once per day for 2 days. For IFNγ injection, mice received 4 consecutive intraperitoneal injections of 400μg/kg BW IFNγ or PBS, once per day for 4 days. For OPZ injection, mice were injected with 150mg/kg BW OPZ or olive oil intraperitoneally, once per week for two weeks. For $CCl_4$ mouse model, mice received 1.6g/kg BW $CCl_4$ intraperitoneally twice per week for 6 weeks. For Western diet mouse model, mice were injected with control or AAV8-ECM1 (VectorBuilder) 7 days before feeding 12-week Western diet (42% fat, 43% carbohydrate, 15% protein) or chow diet. The mice were injected with 100μl virus containing $1.25 \times 10^{11}$ AAV8 vector genomes through the tail vein.

All animal protocols were carried out in full accordance with animal care guidelines and were approved by the local animal care committee, Regierungspräsidium Karlsruhe, Abteilung 3 - Landwirtschaft, Ländlicher Raum, Veterinär- und Lebensmittelwesen (35-9185.81/G-144/22).

**Reagents**

Mouse recombinant EGF (354001), HGF (2207-HG-025) and IFNγ (315-05) were purchased from Corning, R&D and PeproTech, respectively. Human recombinant EGF (AF-100-15) and HGF (294-HG-005) were from PeproTech and R&D systems. Erlotinib (5083S), Oltipraz (sc-205777), hydrogen peroxide (1072102500) were from Cell signaling, Santa Cruz, and Sigma, respectively. The primers, antibodies, siRNA, plasmids and reagent kits used in this study were summarized in **Supplementary Table 1-4**.

**Primary hepatocyte isolation**

8 to 10 weeks old male C57BL/6J mice were used for hepatocyte isolation as previous description (1). The mouse was anesthetized with an intraperitoneal injection of 10%

ketamine hydrochloride (5mg/100mg body weight) and 2% xylazine hydrochloride (1mg/100mg body weight). The mouse liver was sequentially perfused with 50ml perfusion buffer (Krebs-Henseleit buffer with 0.5mM EDTA, Sigma) and 50ml collagenase A buffer (Krebs-Henseleit buffer with 0.1mM $CaCl_2$ and 0.4mg/ml collagenase A, Sigma) through vena cava (at the same time cut portal vein). The perfused liver was removed and minced using forceps to release hepatocytes in suspension buffer. After pipetting cells into 100µm cell strainer, the cell suspension was centrifuged at 50g, 4°C for 5min. To remove dead cells, the supernatant was discarded, Percoll solution (in Hank's buffer) was added and then centrifuged at 270g, 4°C for 10min. The bottom cell pellet was resuspended in culture medium for seeding.

Primary human hepatocytes were isolated by the Department of General, Visceral, Vascular, and Pediatric Surgery, University of Saarland Medical Center, Homburg, using a two-step collagenase perfusion technique with modifications (2). The study protocol complied with national laws and was approved by the local Ethics Committee (approval Nr. 143/21).

**Collagen sandwich hepatocyte culture**

To maintain hepatocellular polarity *in vitro*, we cultured the isolated hepatocytes in collagen sandwich as previously described (3). Briefly, $4 \times 10^5$ freshly isolated hepatocytes were seeded in 6-well plates pre-coated with Rat Tail Collagen I (11179179001, Roche, 250µg/mL in 0.2% acetic acid). After overnight culture, hepatocytes were overlaid with collagen (0.83 mg/mL in 0.2% acetic acid) and cultured in a humidified 37°C incubator with 5% $CO_2$. Hepatocytes were cultured in William's E medium (A1217601, ThermoFisher) supplemented with 10% FBS, 2mM L-glutamine, 1% P/S, 40ng/ml dexamethasone and 0.5% ITS for further experiments.

**Cell culture and treatment**

AML12 cells were grown in DMEM/F-12 medium (21331-020, Gibco) supplemented with 10% Fatal bovine serum (FBS), 2mM L-glutamine, penicillin (100U/mL)-streptomycin (100µg/mL) (P/S), 0.5% Insulin-Transferrin-Selenium (ITS) and 40ng/ml dexamethasone. HEK293T cells were cultured in DMEM medium (11965092, Life

Technologies) supplemented with 10% Fatal bovine serum (FBS), penicillin (100U/mL)-streptomycin (100μg/mL) (P/S). All cells were cultured in the 37°C incubator with a humidified atmosphere containing 5% $CO_2$.

In growth factor/cytokine treatment, cells underwent a 4-to-6-hour starvation in FBS-free medium before the treatment with 100ng/ml epidermal growth factor (EGF), 100ng/ml hepatocyte growth factor (HGF) or 100ng/ml interferon gamma (IFNγ). In the experiments where EGF and/or IFNγ were treated, cells were collected after 24hrs of incubation. For inhibitor experiment, cells were pretreated with erlotinib (1μM) for 2hrs prior to incubation with EGF for 24hrs. $H_2O_2$ (200μM) was treated for 72hrs. For OPZ (50μM), in the co-treatment experiment with EGF, cells were treated with both for 48hrs.

**RNA isolation and qRT-PCR**

All reagents and containers used for RNA extraction were RNase-free grade or treated with 0.1% DEPC (4387937, Thermo Fisher) to eliminate RNase contaminants. Total RNA was isolated from liver tissues or cultured cells using TRIzol (15596018, Invitrogen) according to the manufacturer's instructions. Random primers (SO142, Thermo Fisher), RevertAid H Minus Reverse Transcriptase (EP0452, Thermo Fisher) and RiboLock Rnase Inhibitor (EO0382, Thermo Fisher) were used to reverse transcribe 500ng RNA for cDNA synthesis. qRT-PCR assays were performed on a StepOnePlus system (Applied Biosystem) using SYBR Green Master Kit. The relative quantification of target genes was normalized to the house keeping gene PPIA. Three biological replicates of each condition were measured. The relative fold change in abundance of each target gene compared to a set of internal controls was determined by the $-2^{\Delta\Delta CT}$ formula (4). Primers for qRT-PCR were listed in **Supplementary Table 1**.

**Western Blotting**

Liver tissues or cultured cells were lysed in RIPA buffer (1% Triton X-100, 50mM Tris [pH 7.5], 300mM NaCl, 1mM EGTA, 1mM EDTA and 0.1% SDS) supplemented with protease inhibitors (36978, Thermo Fisher) and phosphatase inhibitors (P5726, Sigma-

Aldrich) on ice for 10min. The supernatant lysates were collected after centrifugation at 13000rpm for 15min at 4°C. Protein concentrations were measured with the Bio-Rad protein assay kit and quantified by absorbance measurements at 562nm via Tecan Infinite M200. After adding LDS sample buffer (4x, NP0007, Lifetechnologies), the samples were boiled at 99°C for 10min. 30μg proteins were separated by SDS-PAGE (8-12% gel) and transferred to 0.2μm Nitrocellulose membranes (10600001, GE Healthcare Life Sciences). After blocking with 5% skimmed milk in TBST (Tris-buffered saline with 0.05% Tween 20) for 1hr at room temperature (RT), the membranes were incubated with primary antibodies overnight at 4°C. The next day, following the TBST washing steps, the membranes were incubated with secondary antibodies for 1hr at RT. After washing with TBST three times, signals were visualized by the Western Lightning Plus-ECL (NEL103001EA, Perkin Elmer) and recorded by the imaging system Fusion SL4 (PEQLAB, Germany). Antibodies used were listed in **Supplementary Table 2**. Each Western blotting experiment was repeated at least three times.

**Nuclear and cytoplasmic protein extraction**

Cells were harvested with trypsin-EDTA and centrifuged at 500g for 5min. The extraction was performed using NE-PER Nuclear and Cytoplasmic Extraction Reagents (78835, Thermo Fisher) according to the manufacturer's instructions.

**RNA interference**

Cells were seeded 24hrs before transfection and expected to be 60-80% confluent when transfection started. Small interfering RNAs (siRNAs) were transfected into cells using Lipofectamine RNAiMAX reagent according to the manufacturer’s instruction. The transfected cells were incubated at 37°C and subjected to different treatments for 24hrs after 24hrs of transfection (30nM each for siEgfr, siErk1, siErk2 and siStat1). siRNAs used in the study were listed in **Supplementary Table 3**. The AllStars Negative Control siRNA (SI03650318, Qiagen) was a negative control.

**Plasmid transfection**

eGFP-STAT1-WT plasmid and eGFP-pcDNA3.1 vector were obtained from Addgene

(Cambridge, MA, USA). Cells were seeded 24hrs before transfection and expected to be 70-80% confluent when transfection started. AML12 were transfected with plasmids using Lipofectamine 3000 (Invitrogen) according to the manufacturer's instruction. The transfected cells were incubated at 37°C for 48hrs (2.5μg per 6-well plate). Plasmids used in the study were listed in **Supplementary Table 3**. The eGFP-pcDNA3.1 vector was a negative control.

**Immunofluorescence staining**

Fresh liver blocks were frozen under dry ice in O.C.T compound (TTEK, Hartenstein) and stored at -80°C. 6μm thick sections were cut by cryotome. Cultured cells were seeded onto 12-well plates with microscope cover glasses (0111580, MARIENFELD) for corresponding experiments. Both tissue slides and cultured cells were fixed on sections with 4% PFA at RT for 15min. After rinsing with PBS three times, slides were permeabilized and blocked with 0.5% Triton X-100 and 1% BSA in PBS for 1hr at RT. Subsequently, the slides were incubated with primary antibodies diluted in PBS with 1% BSA at 4°C overnight. The next day, the slides were washed in PBS three times and then incubated with fluorochrome-conjugated secondary antibodies and DRAQ5 diluted in PBS for 1hr protected from light. After three washes with PBS, slides were mounted with a drop of Fluoromount-G mounting medium (00-4958-02, Invitrogen). Images were scanned under the Leica Microscope Confocal TCS SP8. Antibodies used were listed in **Supplementary Table 2**.

**Immunohistochemistry staining**

4μm paraffin-embedded human liver tissue sections/microarray were deparaffinized and rehydrated. For hematoxylin and eosin (H&E) staining, sections were stained with hematoxylin to visualize nuclei, followed by eosin staining to highlight cytoplasmic components. For Picrosirius Red (PSR) staining, rehydrated sections were incubated with Picrosirius Red solution (0.1% Sirius Red in saturated picric acid) for 1hr to stain collagen fibers. For Nrf2 and α-SMA staining, slides were immersed in citrate buffer PH 6.0 and Tris EDTA PH 9.0, respectively, and boiled for 15min for antigen retrieval. Following cooling at room temperature for 30min, the slides were incubated in blocking peroxide (S200389-2; Dako) for 45min to reduce nonspecific staining. After rinsing in PBS, the tissue slides were incubated with primary antibodies at 4°C overnight. Secondary antibody was added and incubated for 1hr at room temperature. After rinsing in PBS, 3,5-diaminobenzidine (DAB) was used for colour development followed by hematoxylin counterstaining. Finally, all sections were dehydrated and mounted with malinol mounting medium (C9368; Sigma-Aldrich). Slides were scanned using a microscope (DMRBE; Leica) at 20x magnification. Antibodies used were listed in **Supplementary Table 2**.

**Luciferase reporter assay**

Mouse genetic *Ecm1* promoter fragments (-339 ~ +161bp, -953 ~ -454bp) were cloned into pGL4.14 vector (E6691, Promega) and pGL4.13 vector (E6681, Promega), respectively, by Gibson Assembly kit (A46624, Thermo Fisher). Primer sequences are listed in **Supplementary Table 1**. In brief, HEK293T cells were transfected with pGL4.14-Ecm1/pGL4.13-Ecm1 plasmids using lipofectamine 3000 reagents. For luciferase reporter assay, the pGL4.74 plasmid (E6921, Promega) was used as a loading control to standardize the dual-luciferase activity. The cells were co-transfected with STAT1/Nrf2 plasmid for 48 hours. Dual-Luciferase Assay Kit (E2920, Promega) was used to measure the dual-luciferase activity according to the manufacturer's instruction.

**Reactive oxygen species (ROS) detection**

H2DCFDA (Sigma, D399) was dissolved in DMSO for 10mM stock solution. Cultured cells were seeded onto 12-well plates with microscope cover glasses or 96-well plates for corresponding experiments. After treatment, H2DCFDA was added to cells for a final working concentration of 10μM. Following 60min of incubation, ROS-generated fluorescence was detected using a Leica Microscope Confocal TCS SP8 or Tecan Infinite 200 with an excitation wavelength of 488nm and an emission wavelength of 525nm.

**Chromatin immunoprecipitation**

Chromatin immunoprecipitation was performed as previously described with minor modifications (5). For each ChIP, approximately $1\times10^7$ cells were seeded on a 10cm cell culture dish (95% confluence) and treated accordingly. Cells were incubated with 1% PFA for 10min at 37°C to crosslink chromatin and protein. After removing PFA, cells were washed with cold PBS twice and scraped in cold PBS following centrifugation at 1000g, 4°C for 5min. The supernatant was discarded, the cell pellet was resuspended in 500μl lysis buffer (1% SDS, 10mM EDTA, 50mM Tris-HCl pH 8.0) containing 1% protease inhibitor cocktail and lysed on ice for 10min. The chromatin was sonicated using a BioRuptor water bath sonicator (Diagenode), 30sec on/30sec off for 35-40 cycles, to obtain DNA fragments averaging 300-500bp in length. Subsequently, supernatant was collected by centrifugation at 8000g, 4°C for 10min. Remove 50μl from each sonicated sample to serve as input. The remaining chromatin was diluted 1:10 with dilution buffer (0.01% SDS, 1% Triton X-100, 1.2mM EDTA, 16.7mM Tris-HCl pH 8.0, 167mM NaCl). Samples were incubated with 60μl Protein A/G Plus Agarose beads (Santa Cruz), 4μg single-stranded herring sperm DNA on a rotating device for 1h at 4°C. After centrifugation at 2000g, 4°C for 5min, the supernatant was incubated with 5μg primary antibody or IgG for at least 8hrs at 4°C. Subsequently, 60μl Protein A/G Plus Agarose beads and 4μg single-stranded herring sperm DNA were added to samples, and incubated on a rotating device overnight at 4°C. After centrifugation at 2000rpm for 5min, the supernatant was removed, the bottom beads were washed in several buffers with rotation sequentially: TSEI (0.1% SDS, 1% Triton X-100, 2mM EDTA, 20mM Tris-HCl pH 8.0, 150mM NaCl), TSEII (0.1% SDS, 1% Triton X-100, 2mM EDTA, 20mM Tris-HCl pH 8.0, 500mM NaCl), Buffer III (0.25M LiCl, 1% NP-40, 1% Deoxycholic acid, 10mM Tris-HCl pH 8.0) and TE buffer (10mM Tris-HCl pH 8.0, 1mM EDTA) for 10min each at 4°C. After centrifugation at 2000g for 1min, the beads were eluted with 120μl elution buffer (1% SDS, 100mM $NaHCO_3$) for 15min at 30°C. After adding 4.8μL of 5M NaCl and 2μL RNase A (EN0531, Thermo Fisher), the immunoprecipitated complexes together with the inputs were reverse crosslinked at 65°C overnight. Next day, samples were added 2μL proteinase K (EO0491, Thermo Fisher) and incubated while shaking at 60°C for 1h, and then purified with a MinElute PCR Purification Kit, and the DNA obtained was used for subsequent PCR analysis using Phusion High-Fidelity DNA Polymerase and for qRT-PCR analysis using SYBR Green Master Kit. Primers for ChIP were listed in **Supplementary Table 1**. The PCR amplified products were shown on 2% agarose gel electrophoresis.

**Single-cell RNA-sequencing**

The scRNA-seq datasets GSE174748, GSE193850, GSE232182 and GSE202379 were analyzed. Seurat objects were generated from the scRNA-seq gene expression matrix using the Seurat package in R (version 4.4.0). The top 2000 variable genes in each cell scRNA-seq data were identified and normalized using Find Variable Features, Scale Data, and Run PCA functions, which helped to determine the best principal components. Dimensionality reduction was performed using UMAP to effectively summarize these components. Heatmap was utilized to illustrate the expression differences of target genes in hepatocytes between different groups. All clustering and statistical analyses were performed in R (version 4.4.0).

**Gene set variation analysis (GSVA).**

GSVA is an unsupervised, non-parametric method for evaluating transcriptomic gene set enrichment(6). The R package "rWikiPathways" (7) was employed to retrieve a comprehensive collection of 938 pathways and gene sets as reference data. To assess pathway enrichment in advanced and mild liver fibrosis samples from the GSE49541 dataset, the R package "GSVA" was utilized to score the WikiPathways pathways (6).

**Statistical analysis**

Statistical analyses were performed using GraphPad Prism version 6.0 software. Unpaired two tailed Student's t test was used to compare the means between two different groups. One-Way ANOVA was used to determine statistical significance among three or more groups. Variables were described as means and standard deviations (SD). *P* values less than 0.05 were considered statistically significant and indicated as follows: *, $P<0.05$; **, $P<0.01$; and ***, $P<0.001$.

**Supplementary Table 1.** Primers used in the study

**qRT-PCR-Primer**

| Gene | Forward primers (5'-3') | Reverse primers (5'-3') |
|---|---|---|
| *Ppia* | GAGCTGTTTGCAGACAAAGTT | CCCTGGCACATGAATCCTGG |
| *Ecm1* | GCCAGCTCTGTGGAAGTGGA | CCGGAATCTGTTTATGCTTGC |
| *Egfr* | GCCATCTGGGCCAAAGATACC | GTCTTCGCATGAATAGGCCAAT |
| *Stat1* | TCACAGTGGTTCGAGCTTCAG | GCAAACGAGACATCATAGGCA |
| *Nrf2* | CTGAACTCCTGGACGGGACTA | CGGTGGGTCTCCGTAAATGG |
| *Nox4* | AAAGCAAGACTCTACACATCACAT | AGTTGAGGGCATTCACCAAG |
| *Erk1* | TCCGCCATGAGAATGTTATAGGC | GGTGGTGTTGATAAGCAGATTGG |
| *Erk2* | GGTTGTTCCCAAATGCTGACT | CAACTTCAATCCTCTTGTGAGGG |
| *Fos* | CGGGTTTCAACGCCGACTA | TTGGCACTAGAGACGGACAGA |
| *Jun* | CCTTCTACGACGATGCCCTC | GGTTCAAGGTCATGCTCTGTTT |
| *cMyc* | ATGCCCCTCAACGTGAACTTC | CGCAACATAGGATGGAGAGCA |
| *Acta2* | GTCCCAGACATCAGGGAGTAA | TCGGATACTTCAGCGTCAGGA |
| *Timp1* | GCAACTCGGACCTGGTCATAA | CGGCCCGTGATGAGAAACT |
| *Mmp9* | CTGGACAGCCAGACACTAAAG | CTCGCGGCAAGTCTTCAGAG |
| *Col1a1* | GCTCCTCTTAGGGGCCACT | CCACGTCTCACCATTGGGG |
| *Nos2* | GTTCTCAGCCCAACAATACAAGA | GTGGACGGGTCGATGTCAC |
| *Cybb* | TGTGGTTGGGGCTGAATGTC | CTGAGAAAGGAGAGCAGATTTCG |
| *Nqo1* | AGGATGGGAGGTACTCGAATC | AGGCGTCCTTCCTTATATGCTA |
| *Hmox1* | AAGCCGAGAATGCTGAGTTCA | GCCGTGTAGATATGGTACAAGGA |
| *PPIA* | AGCATGTGGTGTTTGGCAAA | TCGAGTTGTCCACAGTCAGC |
| *ECM1* | TGAACCAAATCTGCCTTCCTAAC | GCTGGACTGTGGTAGGTTCCA |

**ChIP-Primer**

| Gene | Forward primers (5'-3') | Reverse primers (5'-3') |
|---|---|---|
| *Ecm1* (-155 ~ +120bp) | GTGCTCCCTTCCATCACCTC | GGCAAGAACTGGTCACTGGT |
| *Ecm1* (-865 ~ -586bp) | GCTCTGCTCCTACCTTTCCTAA | TCACCAATGGACATGACTCAGA |
| *Ppia* | GAGCTGTTTGCAGACAAAGTT | CCCTGGCACATGAATCCTGG |

**Luciferase-Primer**

| pGL4.14 | Primers sequence (5'-3') |
|---|---|
| *Ecm1* P-339 F | GAGCTCTTACGCGTGCTAGCTGAGGCCTCTGACCAGCAAG |
| *Ecm1* P+161 R | AGTACCGGAATGCCAAGCTTGGCCAAGATCAAGGCTGCTCT |
| pGL4.13 | Primers sequence (5'-3') |
| *Ecm1* P-953 F | gtaccgagctcttacgcgtgctagcGTAAAGCCCGGCCCACTTC |
| *Ecm1* P-454 R | tgagatgcagatcgcagatctcgagTTCCTTTCTGTGGGGAGCCCC |

**Supplementary Table 2.** Antibodies used in the study

**Immunoblotting**

| Antibody | Species | Dilution | Company | Cat.No. |
|---|---|---|---|---|
| ECM1 | Rabbit | 1:1000 | Abcam | ab253185 |
| pEGFR Y1068 | Rabbit | 1:1000 | Cell Signaling | 3777T |
| EGFR | Rabbit | 1:1000 | Cell Signaling | 2232S |
| pSTAT1 S727 | Rabbit | 1:1000 | Cell Signaling | 8826S |
| pSTAT1 Y701 | Rabbit | 1:1000 | Cell Signaling | 9167S |

| | | | | |
|---|---|---|---|---|
| STAT1 | Rabbit | 1:1000 | Cell Signaling | 9172T |
| NRF2 | Rabbit | 1:500 | Invitrogen | PA5-27882 |
| p-ERK T202/Y204 | Mouse | 1:1000 | Santa Cruz | sc-7383 |
| ERK 1/2 | Mouse | 1:1000 | Santa Cruz | sc-135900 |
| α-TUBULIN | Rabbit | 1:2000 | Abcam | ab4074 |
| β-ACTIN | Mouse | 1:2000 | Santa Cruz | sc-47778 |
| GAPDH | Mouse | 1:2000 | Santa Cruz | sc-32233 |
| Histone H3 | Mouse | 1:500 | Santa Cruz | sc-517576 |
| Anti-Rabbit IgG | Mouse | 1:5000 | Santa Cruz | sc-2357 |
| Anti-Mouse IgG | Goat | 1:5000 | Santa Cruz | sc-2005 |

**IF Staining**

| Antibody | Species | Dilution | Company | Cat.No. |
|---|---|---|---|---|
| ECM1 | Rabbit | 1:100 | Kindly provided by Prof. Bing Sun | |
| NRF2 | Rabbit | 1:100 | Invitrogen | PA5-27882 |
| EGFR | Rabbit | 1:100 | Cell Signaling | 2232S |
| α-SMA | Mouse | 1:100 | Abcam | ab202368 |
| Col1a1 | Rabbit | 1:100 | Cell Signaling | 72026S |
| LAP-D R58 | Mouse | 1:100 | Cosmo Bio | RIK-MA-R58 |
| DRAQ5 | | 1:1000 | Cell Signaling | 4084L |
| Alexa Fluor 488 Goat anti-Rabbit | Goat | 1:200 | Invitrogen | A-11008 |
| Alexa Fluor 555 Goat anti-Rabbit | Goat | 1:200 | Invitrogen | A-21429 |

**IHC Staining**

| Antibody | Species | Dilution | Company | Cat.No. |
|---|---|---|---|---|
| NRF2 | Rabbit | 1:100 | Invitrogen | PA5-27882 |
| α-SMA | Rabbit | 1:100 | Abcam | ab5694 |
| Goat anti-Rabbit IgG (H+L) Secondary Antibody, HRP | Goat | 1:200 | Invitrogen | 31460 |

**ChIP assay**

| Antibody | Species | Dilution | Company | Cat.No. |
|---|---|---|---|---|
| p-STAT1 S727 | Rabbit | 1:50 | Cell Signaling | 8826S |
| p-STAT1 Y701 | Rabbit | 1:50 | Cell Signaling | 9167S |
| NRF2 | Rabbit | 1:50 | Invitrogen | PA5-27882 |
| IgG Control | Rabbit | 2.5μg/ml | Cell Signaling | 3900S |

**Supplementary Table 3.** siRNA, plasmids and AAV8 used in the study

| Product | **Company** | **Cat.No.** |
|---|---|---|
| siEgfr | Santa Cruz | sc-29302 |
| siStat1 | Santa Cruz | sc-44124 |
| siFos | Santa Cruz | sc-29222 |
| siJun | Santa Cruz | sc-29224 |
| sicMyc | Santa Cruz | sc-29227 |
| siErk1 | Santa Cruz | sc-29308 |
| siErk2 | Santa Cruz | sc-35336 |
| siNrf2 | Santa Cruz | sc-37049 |
| Control siRNA | Qiagen | SI03650318 |

| Product | **Company** | **Cat.No.** |
|---|---|---|

| | | |
|---|---|---|
| eGFP-STAT1-WT | Addgene | 12301 |
| eGFP-pcDNA3.1 | Addgene | 129020 |
| AAV-CMV-Nrf2 | Addgene | 67636 |
| pcDNA3.1 | ThermoFisher | V79020 |
| pGL4.14[luc2/Hygro] | Promega | E6691 |
| pGL4.13[luc2/SV40] | Promega | E6681 |
| pGL4.74[hRluc/TK] | Promega | E6921 |

| Product | **Company** | **Cat.No.** |
|---|---|---|
| AAV8-ECM1 | VectorBuilder | VB230205-1250yhy |

292 **Supplementary Table 4.** Reagent kits used in the study

**Commercial assays**

| Product | Company | Cat.No. |
|---|---|---|
| SYBR Green Master Kit | Thermo Fisher | A25918 |
| Bio-Rad protein assay kit | Bio Rad | 5000113-115 |
| Lipofectamine® RNAiMAX kit | Thermo Fischer | 13778-075 |
| Lipofectamine™ 3000 Transfection Reagent | Thermo Fischer | L3000008 |
| Phusion® High-Fidelity DNA Polymerase | New England Biolabs | M0530S |
| MinElute PCR Purification Kit | Qiagen | 28006 |
| NE-PER™ Nuclear and cytoplasmic extration reagents | Thermo Fisher | 78835 |
| PureLink Quick Plasmid Miniprep Kit | Invitrogen | K210010 |
| PureLink® HiPure Plasmid Maxiprep Kit | Invitrogen | K210007 |
| Gibson Assembly HiFi Cloning Kit | Thermo Fisher | A46624 |
| Dual-Luciferase Assay Kit | Promega | E2920 |

### Supplementary figures and figure legends

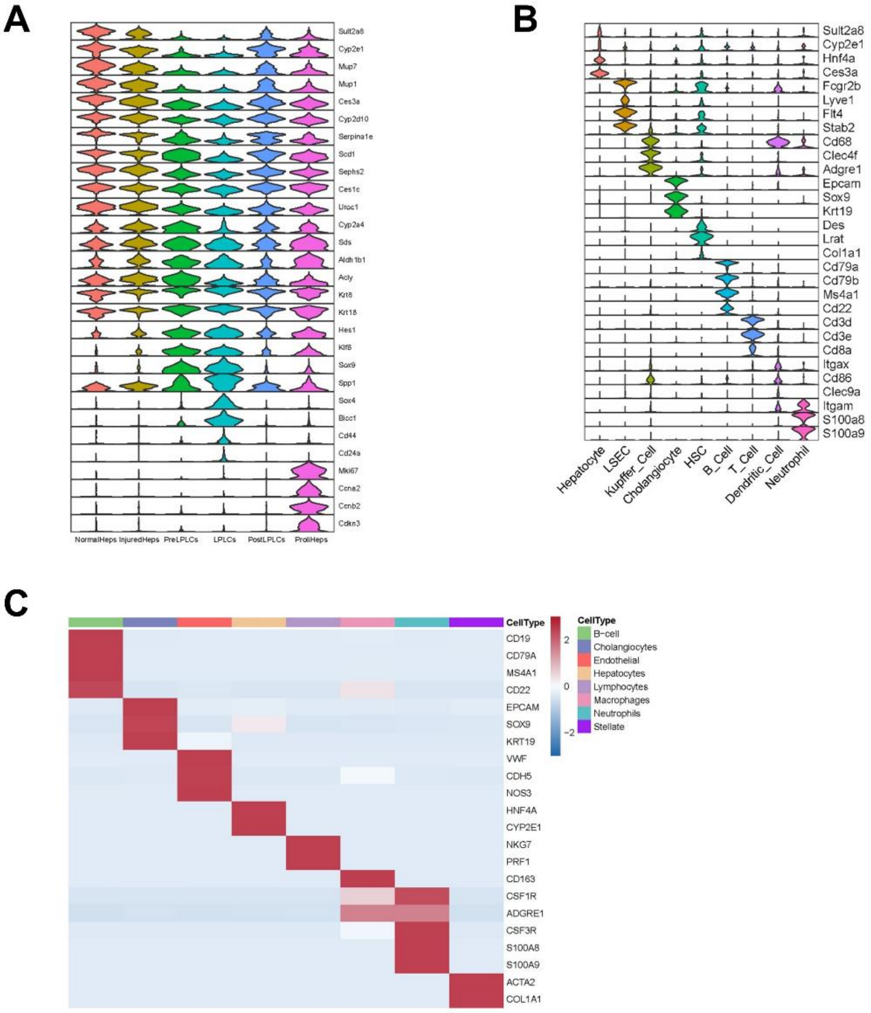


**Suppl. Fig. 1. Single-cell RNA-sequencing analysis of different mouse models and patients with CLD**

Marker genes for defining different cell clusters in **(A)** DDC diet (GSE193850), **(B)** CD-HFD fed (GSE232182) mouse models, and **(C)** patients with CLD (GSE202379).

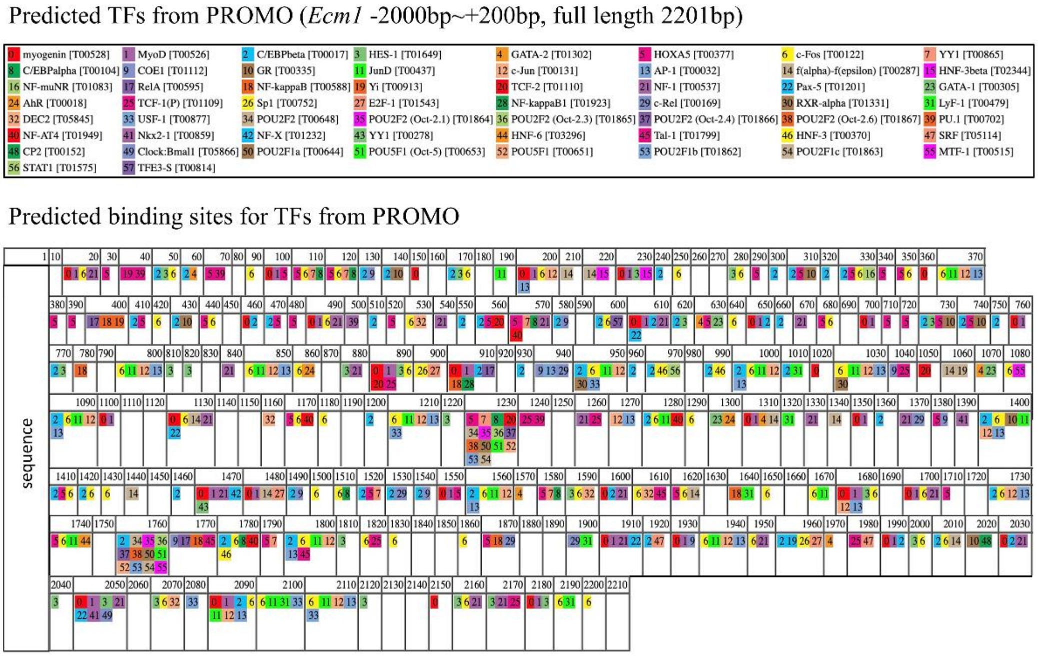


298 **Suppl. Fig. 2. Predictive transcription factors and binding sites in *Ecm1* gene promoter**

Predicted transcription factors and their binding sites for *Ecm1* promoter (-2000 ~ +200bp relative to the TSS of the *Ecm1* gene) generated by PROMO.

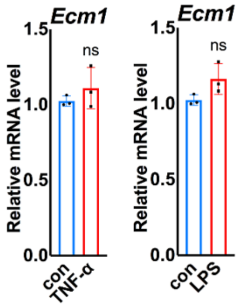


300 **Suppl. Fig. 3. Expression of *Ecm1* in MPH after TNF-α and LPS treatment**

301 qRT-PCR for the impact of TNF-α (4ng/ml) and LPS (5μg/ml) on mRNA expression of *Ecm1* in MPH. The results of qRT-PCR were normalized to *Ppia*. *P*-values were calculated by unpaired Student's t test. Bars represent the mean ± SD. *, *P*<0.05; **, *P*<0.01; ***, *P*<0.001.

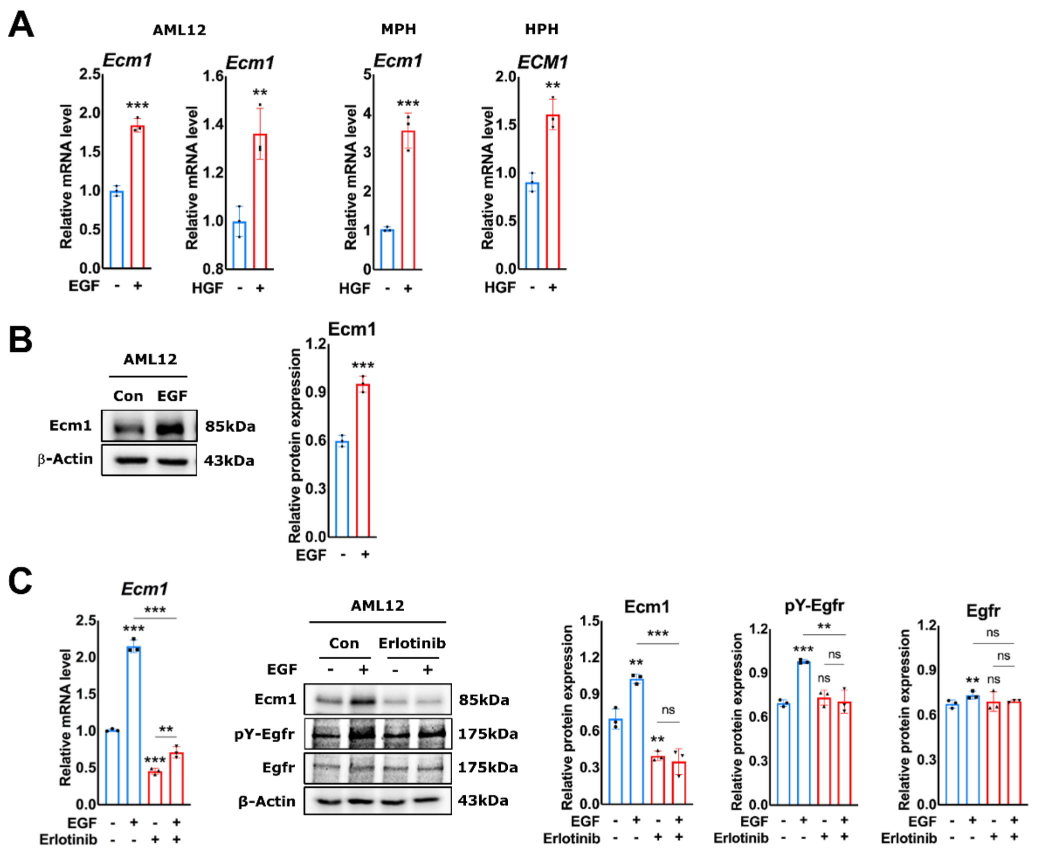


**Suppl. Fig. 4. The effect of growth factors on ECM1 expression in AML12, mouse and human primary hepatocytes**

**(A)** qRT-PCR for *Ecm1/ECM1* mRNA expression in AML12, MPHs and HPHs with or without EGF/HGF treatment for 24hrs. **(B)** Immunoblotting of Ecm1 protein expression in AML12, treated with EGF for 24hrs. **(C)** qRT-PCR and immunoblotting showing the effect of erlotinib on Ecm1 expression and Egfr Y1068 phosphorylation in AML12 with or without EGF treatment. The results of qRT-PCR were normalized to *PPIA*. β-Actin is a loading control. Quantification of protein expression was performed by ImageJ (National Institutes of Health, Bethesda, Maryland, USA). *P*-values were calculated by unpaired Student's t test. Bars represent the mean ± SD. *, $P<0.05$; **, $P<0.01$; ***, $P<0.001$.

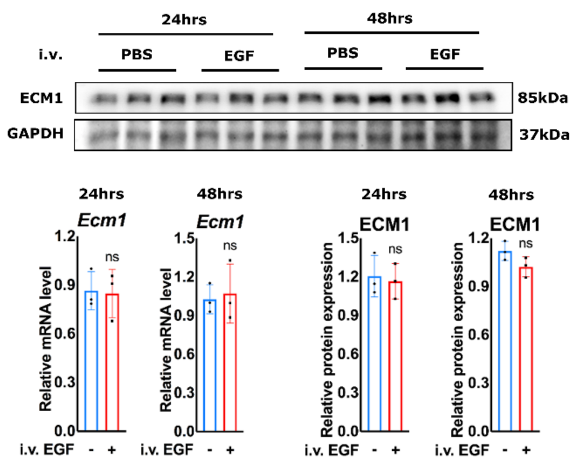


**Suppl. Fig. 5. Effect of EGF on the regulation of Ecm1 expression *in vivo***

qRT-PCR, immunoblotting for Ecm1 expression in liver tissues from mice treated with or without EGF. The results of qRT-PCR were normalized to *Ppia*. Gapdh is a loading control. Quantification of protein expression was performed by ImageJ (National Institutes of Health, Bethesda, Maryland, USA). *P*-values were calculated by unpaired Student's t test. Bars represent the mean ± SD. *, *P*<0.05; **, *P*<0.01; ***, *P*<0.001.

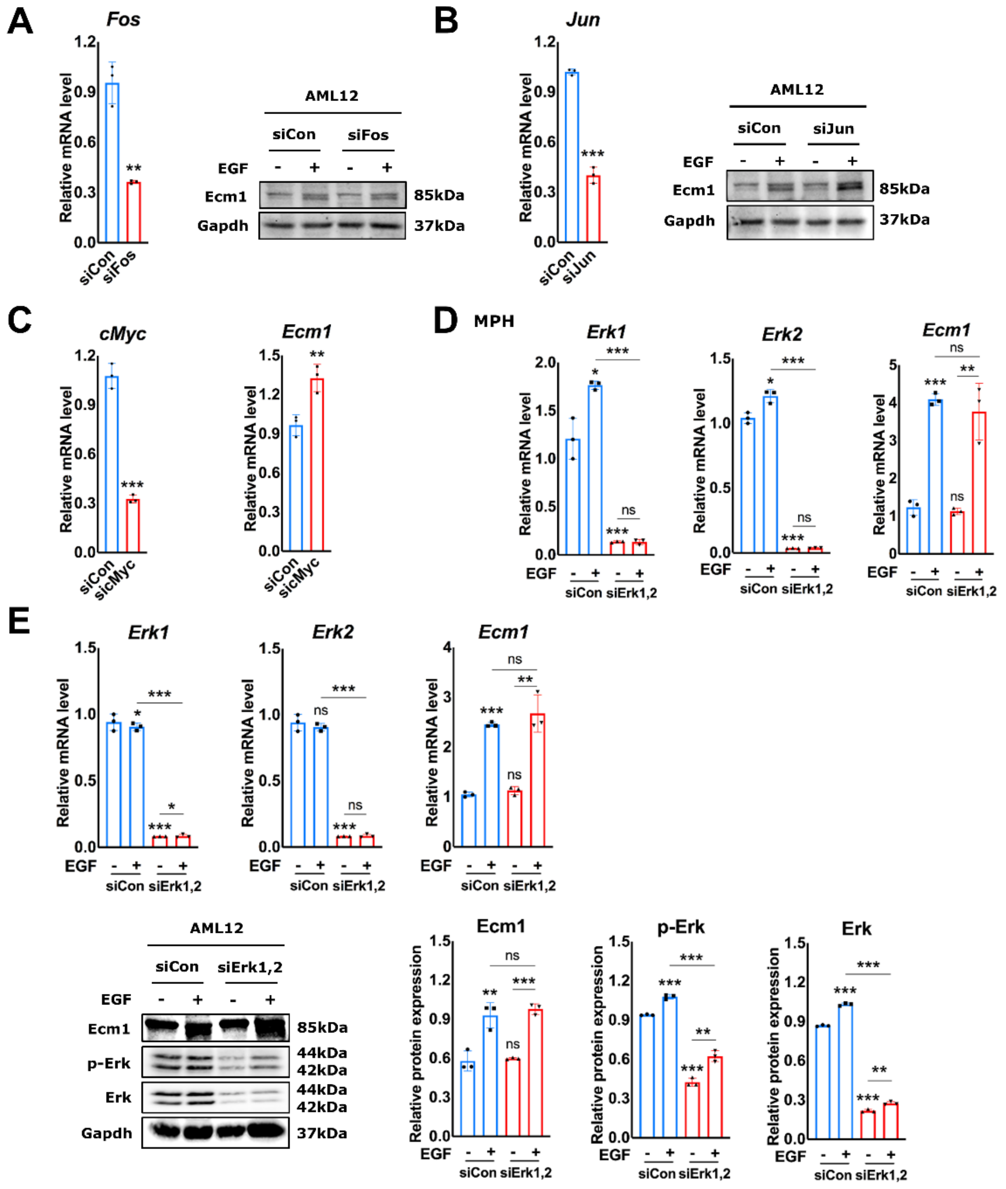


**Suppl. Fig. 6. Downstream signaling pathway of EGF in the regulation of Ecm1 expression in hepatocytes**

Western blotting showing the impact of **(A)** *Fos* and **(B)** *Jun* knockdown on protein expression of Ecm1 in EGF-treated AML12. qRT-PCR showing knockdown efficiency of siRNA. **(C)** qRT-PCR for the impact of *cMyc* knockdown on mRNA expression of *Ecm1*, *cMyc* in AML12. **(D)** qRT-PCR for the impact of *Erk1* and *Erk2* knockdown on mRNA expression of *Ecm1*, *Erk1* and *Erk2* in EGF-treated MPHs. **(E)** qRT-PCR and Western blotting showing the impact of *Erk1* and *Erk2* knockdown on mRNA and protein expression of Ecm1, Erk1 and Erk2 in EGF-treated AML12. Immunoblots for p-Erk T202/Y204 are also shown. The results of qRT-PCR were normalized to *Ppia*. Gapdh is a loading control. Quantification of protein expression was performed by ImageJ (National Institutes of Health, Bethesda, Maryland, USA). *P*-values were calculated by unpaired Student's t test. Bars represent the mean ± SD. *, $P<0.05$; **, $P<0.01$; ***, $P<0.001$.

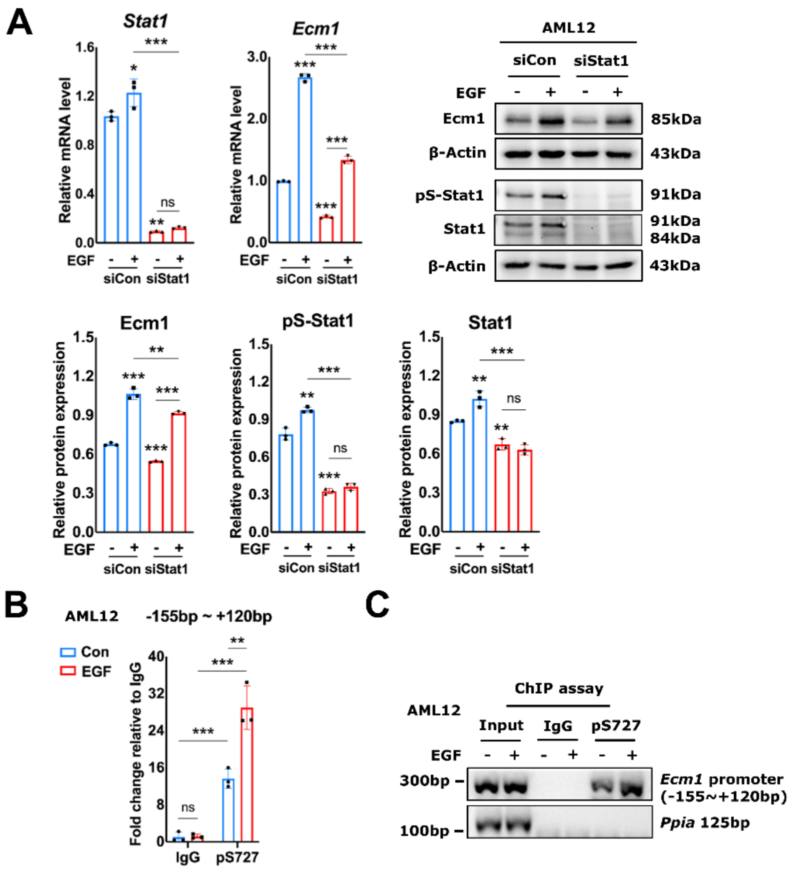


**Suppl. Fig. 7. EGF/EGFR regulates Ecm1 expression through Stat1 in AML12**

**(A)** qRT-PCR and Immunoblot data for mRNA and protein expression of Ecm1 and Stat1 in EGF-treated AML12 with or without RNAi mediated *Stat1* knockdown. Additionally, p-Stat1 S727 levels are shown. The results of qRT-PCR were normalized to *Ppia*. β-Actin is a loading control. Quantification of protein expression was done by ImageJ (National Institutes of Health, Bethesda, Maryland, USA). **(B)** ChIP qRT-PCR showing the binding of p-Stat1 S727 to the *Ecm1* gene promoter in AML12 cells treated with or without EGF for 24hrs. Relative fold change of immunoprecipitated genomic fragments in EGF-treated cells was compared to untreated control group. Fragment "-155bp ~ +120bp" was relative to the transcription start site of *Ecm1* gene. Rabbit IgG-bound chromatin served as a negative control. **(C)** The PCR amplified products of "-155bp ~ +120bp" fragments were shown on a 2% agarose gel. *Ppia* represents non-specific binding. *P*-values were calculated by unpaired Student's t test. Bars represent the mean ± SD. *, *P*<0.05; **, *P*<0.01; ***, *P*<0.001.

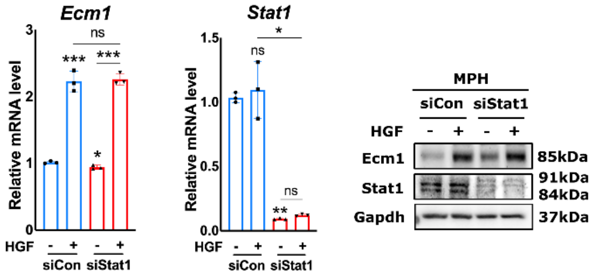


**Suppl. Fig. 8. HGF regulates Ecm1 expression independent of Stat1**

qRT-PCR and Western blotting showing the impact of *Stat1* knockdown on mRNA and protein expression of Ecm1 and Stat1 in HGF-treated MPHs. The results of qRT-PCR were normalized to *Ppia*. Gapdh is a loading control. *P*-values were calculated by unpaired Student's t test. Bars represent the mean ± SD. *, *P*<0.05; **, *P*<0.01; ***, *P*<0.001.

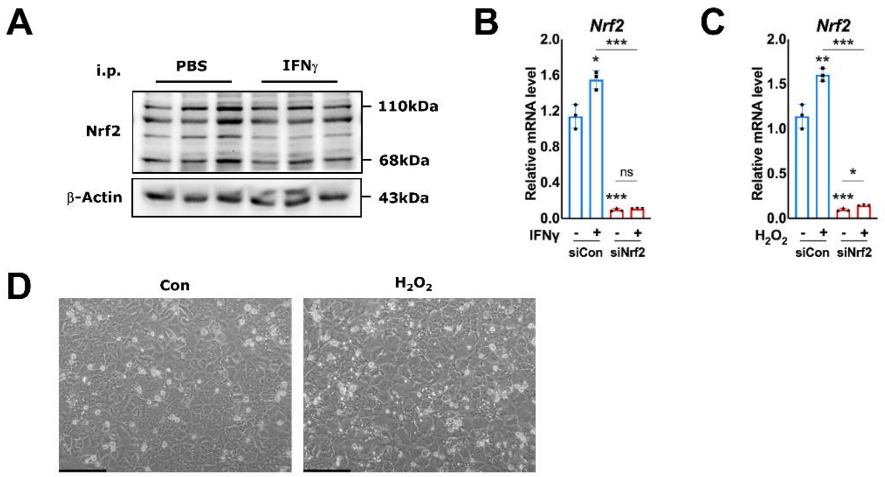


**Suppl. Fig. 9. Nrf2 expression and bright-field images of MPHs**

**(A)** Western blotting showing protein level in the liver tissues from mice treated with PBS or IFNγ (400μg/kg/day, i.p., for 4 days). β-Actin is a loading control. **(B-C)** qRT-PCR showing the knockdown efficiency of siNrf2 in MPHs. The results of qRT-PCR were normalized to *Ppia*. **(D)** Bright-field images of MPHs with or without $H_2O_2$ treatment. Scale bar 200μm. *P*-values were calculated by unpaired Student's t test. Bars represent the mean ± SD. *, *P*<0.05; **, *P*<0.01; ***, *P*<0.001.

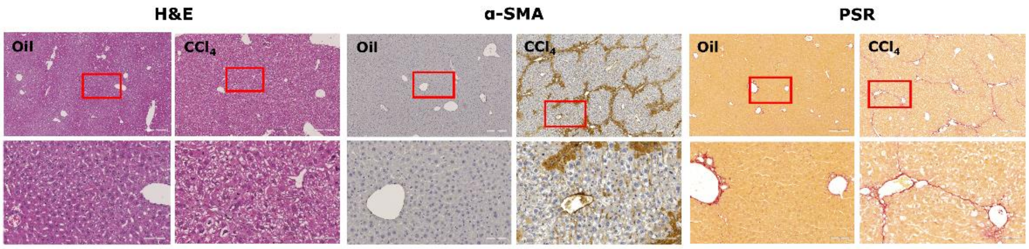


**Suppl. Fig. 10. IHC staining in $CCl_4$-treated mice**

H&E, α-SMA and PSR staining showing liver damages in the mice treated with oil or $CCl_4$ (1.6g/kg BW $CCl_4$, i.p., twice per week).

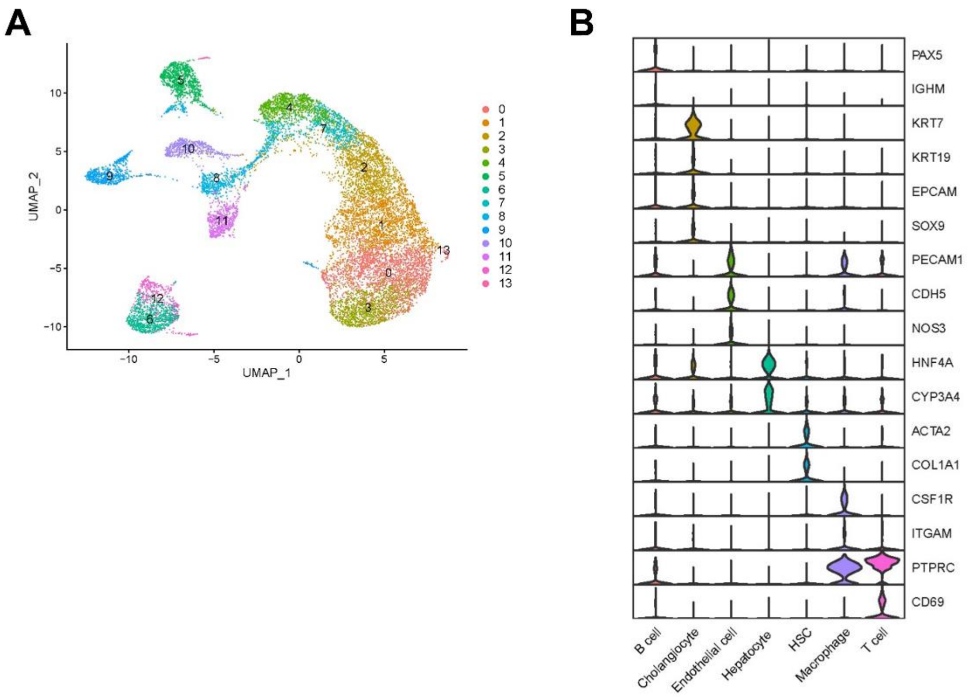


**Suppl. Fig. 11. Single-cell RNA-sequencing analysis of normal livers and cirrhotic livers from MAFLD patients**

**(A)** UMAP visualization of cell populations. **(B)** Marker genes for defining different cell clusters.
